# DeepPheno: A Deep Learning Framework for Linking Hyperspectral Imaging and SNP Genotypes in Lettuce

**DOI:** 10.64898/2026.07.09.737449

**Authors:** Frank G. Okyere, Sarah L. Mehrem, Basten L. Snoek, Guido Van den Ackerveken, Sanne Abeln

## Abstract

While whole-genome sequencing captures millions of single nucleotide polymorphisms (SNPs) and hyperspectral imaging (HSI) enables non-destructive plant phenotyping, integrating these modalities to link genotype to phenotype remains challenging due to their high dimensionality and non-linearity. This study presents DeepPheno a deep learning framework that predicts SNP genotypes from HSI data, using model predictability as a proxy for genotype-phenotype association. HSI data were acquired from 194 lettuce genotypes under field conditions. HSI data patches (20×20 pixels × 224 spectral bands) were used to train a hybrid CNN to predict the variant of a specific SNP. The framework was validated on SNPs with known phenotypic effects (anthocyanin, leaf serration, pale pigmentation), achieving high predictive performance (AUC ranging from 0.806 to 0.935), whereas models trained on randomly shuffled labels performed at chance (mean AUC ≈ 0.51). Extending the workflow to 50 randomly selected putatively neutral SNPs, most yielded low predictability, but two showed high performance (AUC > 0.76), suggesting uncharacterized genotype-phenotype links. Explainable AI, including SHAP and Grad-CAM, identified relevant spectral and spatial features driving these predictions, particularly the green and red-edge wavelengths associated with pigment dynamics and leaf structure. These results establish a framework for understanding complex genotype-phenotype interactions in plants and extracting these links from HSI data without predefining the exact trait values. It provides an avenue for high-throughput trait discovery and description and extends the integration of image-based phenomics with plant genetics.

## 1. INTRODUCTION

Advances in crop productivity, resilience, and nutritional quality are essential to meet the projected doubling of global food demand by 2050 [1]. Achieving this goal requires accelerating plant breeding, which fundamentally depends on understanding the relationship between genetic variation and phenotypic traits that can be targeted for crop improvement. Recent advances in next-generation sequencing technologies have enabled the identification of millions of single-nucleotide polymorphisms (SNPs), providing a scalable resource for studying genetic variation in plants[2] While many SNPs are phenotypically neutral, functional variants can influence important traits such as anthocyanin biosynthesis[2] and leaf serration[3] in lettuce. However, translating genomic information into robust phenotypic understanding remains a major challenge in plant phenomics, largely due to limitations in phenotyping approaches[4]. Traditional phenotyping methods are often destructive, labor-intensive, and time-consuming, thereby restricting their application across large breeding populations[5].

To address these limitations, advanced high-throughput phenotyping methods are required. Hyperspectral imaging (HSI) has emerged as a powerful alternative, enabling non-destructive and high-throughput measurement of plant biochemical and physiological properties. By integrating imaging and spectroscopy, HSI captures contiguous narrow-band spectral information across plant growth stages, allowing detailed characterization of plant physiology[6]. In addition, HSI-derived spectral indices have been shown to associate with trait-related genetic variation[7], and the technology enables early detection of stress responses prior to the appearance of visible symptoms[8]. Despite HSI data providing rich phenotypic information, mapping these traits, beyond the individual wavelengths, to underlying genetic variation is a key challenge. Genome-wide association studies (GWAS) have been widely used to identify statistical associations between SNPs and phenotypic traits[9]. The integration of HSI with genetic mapping approaches including GWAS, has successfully uncovered the genetic architecture of complex traits across various crops. These include mapping temporal spectral QTLs in field-grown lettuce[3], evaluating nitrogen response mechanisms in wheat[10] and identifying seed vigor loci in peanut[11]. However, these approaches typically rely on predefined, manually measured traits or use classical RGB image based phenotyping, these may not fully capture the complex, high-dimensional phenotypic variation embedded in hyperspectral data. While HSI does not directly measure genetic variation, it does capture the effect of SNPs with an influence on plant biochemical and physiological processes, and therefore in turn generate detectable spatial-spectral patterns. Consequently, there is a need for computational approaches capable of modeling nonlinear relationships between high-dimensional hyperspectral signals and underlying genetic variation.

Recent studies integrating HSI with machine learning (ML) have demonstrated promising progress in addressing these challenges[12]. Machine learning approaches have been widely applied to plant stress detection[7],trait identification[13], and crop monitoring. For instance, regression models have been used to estimate biochemical traits including chlorophyll content, nitrogen status, and water content from HSI data. Furthermore, deep learning methods have significantly improved predictive performance by capturing complex nonlinear relationships in HSI data [14]. Convolutional neural networks have been employed for early stress detection with higher sensitivity than traditional vegetation indices[15], while deep residual networks have achieved high accuracy in seed variety classification using hyperspectral inputs[16].In addition, feature selection techniques integrated with deep learning models have enhanced the non-destructive estimation of plant growth indicators such as Soil Plant Analysis Development (SPAD) values and biomass[17].

Despite these advances, there remains a lack of systematic frameworks that directly link genetic variants to high-dimensional hyperspectral phenotypes. Most HSI-based approaches focus on predicting phenotypic traits rather than inferring genotypes at the level of individual SNPs, thereby limiting the resolution required to dissect the functional role of specific genetic variants[18]. Similarly, GWAS approaches rely on handcrafted or aggregated traits as proxy phenotypes, which may fail to capture the full richness of hyperspectral data. Another major limitation is the “black-box” nature of many machine learning models, which often lack interpretability and thus provide limited biological insight into how spectral signals relate to underlying physiological or biochemical processes.

These limitations highlight the need for integrative frameworks that can quantify how genetic variation is expressed through measurable spatial-spectral phenotypes. Such frameworks would enable systematic evaluation of whether functional genetic variants produce detectable signatures in hyperspectral data through their influence on plant biochemistry and physiology. Moreover, they would facilitate the identification of specific spectral features associated with genetic variation, thereby enabling the discovery of novel traits and supporting marker-assisted breeding strategies.

Building on this need, this study proposes a complementary perspective: instead of predicting phenotype from genotype, we predict SNP genotypes from HSI phenotypes as a proxy for genotype–phenotype association. We hypothesize that functional SNPs, those linked to variation in a pre-defined trait, alter plant biochemical properties, thereby generating consistent and learnable signatures in hyperspectral data, whereas neutral variants do not produce detectable patterns (Figure 1). Under this framework, the ability of a model to predict the SNP variant from hyperspectral features provides a direct measure of the biological impact of this SNP. High predictive performance indicates a meaningful genotype-phenotype association, while near-random performance suggests limited or no detectable HSI phenotypic effect.

**Figure 1.**
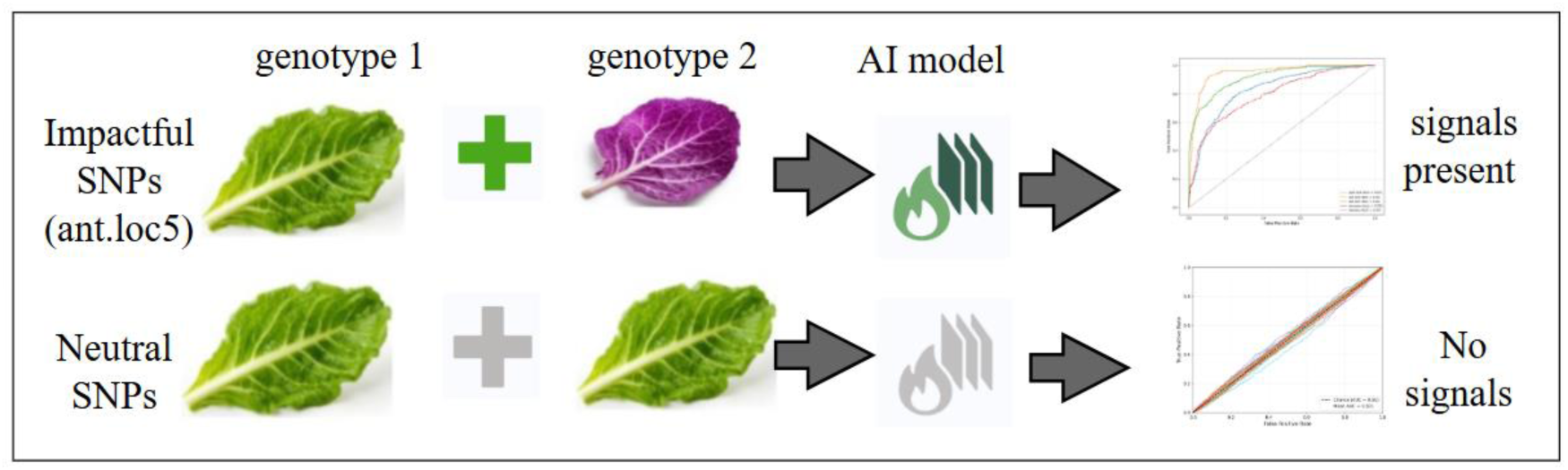
Conceptual framework for detecting genotype–phenotype associations via SNP genotype classification from hyperspectral phenotypes. Functional SNPs trigger biochemical and structural changes that generate measurable spectral reflectance variations, producing learnable signals. In contrast, neutral SNPs do not produce detectable biochemical changes, resulting in near-random model performance.

To test this hypothesis, we present DeepPheno, an integrative deep learning framework that combines HSI, genomic data, and explainable AI (XAI). The framework comprises four key components: (1) HSI preprocessing and spatial-spectral patch extraction; (2) development and validation of deep learning models for SNP genotype prediction; (3) estimation of null distributions through random label shuffling to assess the statistical significance of genotype-phenotype associations; and (4) application of XAI techniques to identify spectral and spatial regions driving model predictions. This framework enables systematic identification of SNPs with measurable phenotypic effects and provides a scalable approach for discovering novel genotype-phenotype associations. Collectively, this work establishes a robust link between plant genetic variation and its resulting high-dimensional spatial-spectral phenotypes, addressing a critical challenge in high-throughput plant phenotyping and genomic selection.

## 2. MATERIALS AND METHODS

In this study, multiple image analysis pipelines were developed to extract features from hyperspectral imaging (HSI) data for SNP genotype prediction. These pipelines were used to assess genotype-linked spectral variation in four SNPs: anthocyanin accumulation (2 SNPs), leaf serration (1 SNP), and pale pigmentation (1 SNP). The overall workflow consists of four main stages (Figure 2): (1) HSI preprocessing, (2) deep learning model design and validation, (3) statistical baseline analysis, and (4) explainable AI-based biological interpretation–including SHAP and GRAD-CAM.

**Figure 2.**
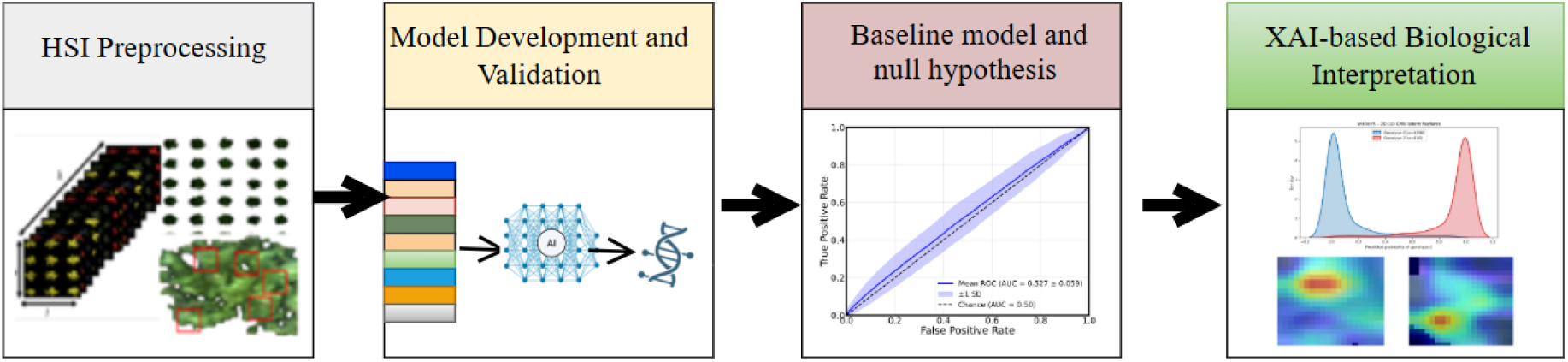
Proposed pipeline for this study. (1)HSI processing to obtain patches, GWAS data analysis to get the **SNPs with known phenotypic impact** (2) Design and validation of ML models to predict SNP status (3) Baseline for statistical analysis for null hypothesis, (4) explainable AI-based biological interpretation.

### 2.1 Plant Materials and Experimental Design

A total of 194 lettuce (*Lactuca sativa*) genotypes were cultivated under field conditions at Maasbree in the Netherlands [3] to capture genetic variation in plant physiological and biochemical traits. Plants were cultivated following standard agronomic practices, and measurements were conducted at defined developmental stages to ensure consistency across samples. The accessions were arranged in a complete randomized block pattern with each plot having about 36 plants. Each plot had a replicate and was assigned a unique identifier, allowing linkage between HSI data and corresponding genotypic information. This experimental design enables integration of phenotypic and genomic datasets at the individual plant level. Three phenotypes with four linked SNPs (anthocyanin levels (purple color; 2 SNPs), serration (1 SNP), and chlorophyll content (pale color; 1 SNP)) were imaged across five developmental time points spanning the full growing season. This study focuses on a cross-sectional analysis using the pooled multi-temporal dataset rather than explicit temporal modelling. Figure 3 shows pseudo-RGB images of plants at 57 DAS and 78 DAS, representing early and late developmental stages respectively.

**Figure 3.**
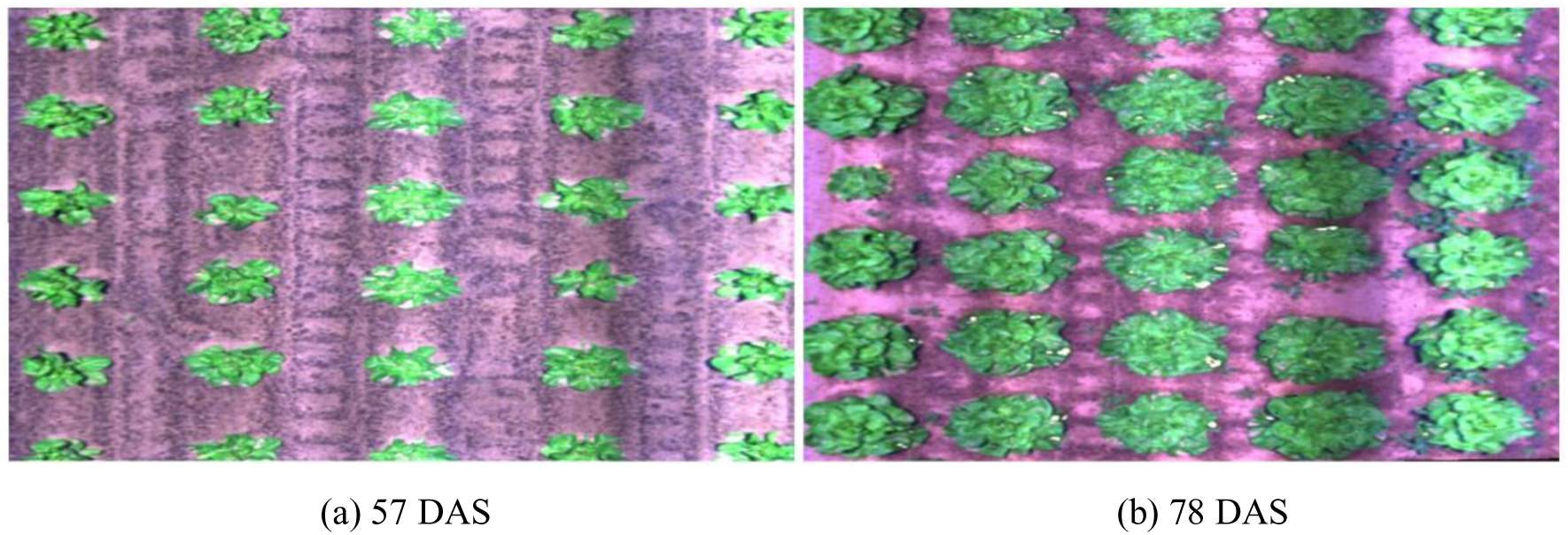
Pseudo RGB images of lettuce accessions grown under field conditions. Panels show representative plots at two time points (a) 57 DAS, when rosettes were smaller and less compact, and (b) 78 DAS, when plants were fully expanded and displayed clearer phenotypic differences. Each image shows over 25 individual plants corresponding to single-accession replicates.

### 2.2 Hyperspectral Image Acquisition and Pre-processing

Hyperspectral images were acquired across multiple developmental stages of lettuce using an FX10 spectral camera (Specim, Finland), covering the spectral range of 400–1000 nm across 224 contiguous bands (also described in Mehrem *et al.* [19]). The camera is a push-broom type operating within the visible and near-infrared (VNIR) regions (390–1024 nm). It captures one row of spatial pixels per time frame, with full spectral information (224 bands) recorded for each pixel. The sensor uses a 5.5 nm full width at half maximum (FWHM) slit and samples data at approximately 2 nm spectral intervals.. Each HSI data was stored as a three-dimensional cube (height × width × spectral bands), capturing both spatial and spectral information.

Raw HSI data were preprocessed following a multi-step pipeline, including radiometric calibration, denoising, normalization, and segmentation. Radiometric calibration was applied to correct for illumination-induced variability arising from changes in solar angle, atmospheric conditions, and cloud cover. Dark and white reference panels were included during scanning, and calibration was performed using standard radiometric correction equations[20]. Spectral smoothing and denoising were applied to improve the signal-to-noise ratio (SNR) by eliminating outliers while preserving spectral features. A Savitzky–Golay filter (a conventional low-pass filter) was used, fitting a second-degree polynomial within a symmetric window across spectral bands. Optimal parameters (window size = 13, polynomial order = 2) were selected to balance noise reduction and feature preservation, avoiding artifacts from smaller windows and over smoothing from larger ones[21].

HSI segmentation was performed using a two-step strategy to remove background elements (soil, stones, and weeds). First, normalized difference vegetation index (NDVI)-based pseudo-labels were generated using 910 nm and 950 nm wavelengths. These binary masks served as proxy ground truth labels to train a random forest segmentation model using non-overlapping 3D patches. Supplementary Figure S1 illustrates the segmentation workflow and representative results.

### 2.3 SNP Genotype Data and Preprocessing

All 194 accessions were genotyped following the workflows of van Workum et al.[22] *D*ijkhuizen and van Eijnatten et al. [3] with data from van Workum *et* al. [22] and Wei et al.[23]. Raw Illumina reads were aligned to the *Lactuca sativa* ‘Salinas’ v8 reference genome, and variants were called using the GATK HaplotypeCaller pipeline. The resulting SNPs were subjected to quality control, including filtering and linkage disequilibrium (LD) clumping, to obtain a set of independent loci (as described in Dijkhuizen et al [1]). Variants were filtered based on genotype call rate (≥90%), minor allele frequency (MAF ≥5%), and Hardy-Weinberg equilibrium to remove low-quality or potential genotype errors.

After quality control, a total of 2,491,009 high-quality SNPs were retained for GWAS and downstream analyses. SNPs were encoded numerically as 0 (homozygous reference), 1 (heterozygous), and 2 (homozygous alternate), enabling their use in downstream analysis. To enhance class separability and reduce ambiguity in phenotype-genotype mapping, heterozygous samples (state = 1) were excluded from downstream analysis. We did this based on the expectation that intermediate phenotypic expression may obscure spectral distinctions between genotype classes. Consequently, all classification tasks were formulated as binary problems To select labels for model training, three SNPs groups were considered: (i) GWAS-identified SNPs with known phenotypic impact (ii) Permutated SNPs that lacks genotype-phenotype association and (iii) randomly selected putatively neutral SNPs serving as controls to evaluate the model’s ability to distinguish biologically impactful genotypes from likely neutral variants.

#### 2.3.1 SNPs with known phenotypic impact

Genome-wide association studies (GWAS) were performed on three physiological traits, anthocyanin pigmentation, leaf serration, and chlorophyll content (pale coloration), using the lme4qtl R-package[24], following the approaches described by Dijkhuizen et al. [3] and Mehrem et al. [25]. Population structure was accounted for by including a SNP-based covariance matrix, and a conservative significance threshold of −log₁₀(P) > 7 was applied to correct for multiple testing. The robustness of the GWAS was validated in two ways: (i) consistency of QTL profiles between clamped and unclamped datasets, and (ii) high concordance across independent experimental replicates. Candidate genes were identified using the *Lactuca sativa* ‘Salinas’ v8 genome annotation.. Four positive control SNPs with known biological effects were selected for downstream analysis based on *Lactuca sativa* V8 genome positions:

1. ant.loc5 (Chr5, 86.7 Mbp): targets the transcription factor *LsMYB114* within the *RLL4* pigmentation locus[2] - for anthocyanin
2. ant.loc9 (Chr9, 95.1 Mbp): anchors the *LsANS/LsLDOX* structural enzyme within the *RLL2* pathway [2] - for anthocyanin
3. pale.loc4 (Chr4, 106.0 Mbp): co-localizes with *LsGLK*, an upstream chloroplast-regulating transcription factor [25] - for pale colour
4. serr.loc5 SNP (Chr5, 114.8 Mbp): targets *LsTCP4*, a transcription factor governing leaf margin morphogenesis[26] - for serration

Hereafter, these loci are referred to as ant.loc5, ant.loc9, pale.loc4, and serr.loc5, respectively, based on their chromosomal locations and associated phenotypes

#### 2.3.2 Null distribution (Random Labeling)

To evaluate whether the model performance reflects true biological signal rather than spurious correlation, a null distribution was constructed using random label permutation. To do this, true genotype labels were randomly shuffled while preserving class balance and sample size. This procedure disrupts genuine genotype-phenotype associations while maintaining the HSI data structure. Label permutation was repeated 300 times to obtain a stable estimate of the null distribution. For each permutation, model performance was evaluated using area under the receiver operating characteristic curve (AUC) and classification accuracy. The aggregated results were used to construct an empirical null distribution plot representing expected model performance under the hypothesis of no genotype-phenotype relationship. Statistical significance was assessed by comparing model performance of the true genotype labels to the null distribution. The 95^th^ percentile of the null distribution was used as the threshold (α = 0.05). p-values were computed as the proportion of permuted runs yielding performance equal to or above that of the true labels. This approach provides a statistical baseline for distinguishing meaningful genotype-phenotype signals from overfitting or high-dimensional artifact.

#### 2.3.3 Selection of randomly sampled putatively neutral control SNPs

To establish a negative control framework, a set of putatively neutral SNPs with no known or expected association with the target phenotypes was curated. These SNPs—hereafter referred to as randomly selected SNPs—serve to determine whether observed genotype–phenotype associations arise from true biological signals in the HSI data rather than from artifacts, population structure, or linkage to known functional variants. Additionally, this approach enables the identification of unexpected predictive signals that could reflect previously uncharacterized genotype–phenotype relationships.

SNP selection followed a controlled pipeline (**Supplementary Figure S2**) designed to match the quality and minor allele frequency (MAF) characteristics of the known phenotypic SNPs while ensuring genetic independence and likely phenotypic neutrality. From the full SNP matrix, quality filtering was applied to retain variants with call rate ≥95% and MAF ≥3%, removing rare variants with limited statistical power and maintaining allelic balance. To avoid linkage with known functional loci, linkage disequilibrium (LD) pruning was performed by excluding: (i) SNPs with pairwise correlation |r| > 0.3 with any of the four known phenotypic SNPs, and (ii) all SNPs within a 1 Mb window on the same chromosome as any of these four SNPs. Furthermore, to mitigate confounding effects due to population structure, SNPs strongly correlated with the first two principal components of genetic variation (variance explained <5%) were excluded. The remaining SNPs were considered putatively neutral relative to the known loci and unlikely to exhibit phenotypic effects linked to causal variants.

From this filtered pool of putatively neutral variants (about 10458 SNPs), 50 SNPs were randomly sampled for downstream analysis. The MAF distribution of the selected SNPs was verified to be comparable to that of the known phenotypic SNPs. Selected SNPs were labeled using a standardized format (e.g., MAF_0.123_chr1_123456), indicating MAF, chromosome, and genomic position.

### 2.4 Deep Learning for SNP Identification

To predict SNP genotypes from hyperspectral imaging (HSI) data, we developed a hybrid convolutional neural network (CNN) architecture that extends the 3D-2D CNN framework of Okyere et al [2] by incorporating an additional spectral feature extraction branch.

#### 2.4.1 Proposed Model Architecture

The baseline 3D-2D CNN model extracts joint spatial-spectral features using 3D convolutions, followed by reshaping and 2D convolutions to extract higher-level spatial representations. While effective, this approach limits detailed spectral feature extraction due to the computational expense of the 3D convolutional block, which limits kernel usage. To address this limitation, the architecture was modified by channeling the feature maps of the 3D block into two parallel branches: 2D and 1D CNN blocks (Figure 4). The spatial branch receives the reshaped feature maps from the 3D block and applies three homogenous depthwise separable convolutional layers, each followed max-pooling and batch normalization layers. This branch captures structural and spatial patterns such as leaf shape, venation, and surface texture, enabling efficient and dense spatial feature learning while maintaining computational efficiency.

**Figure 4.**
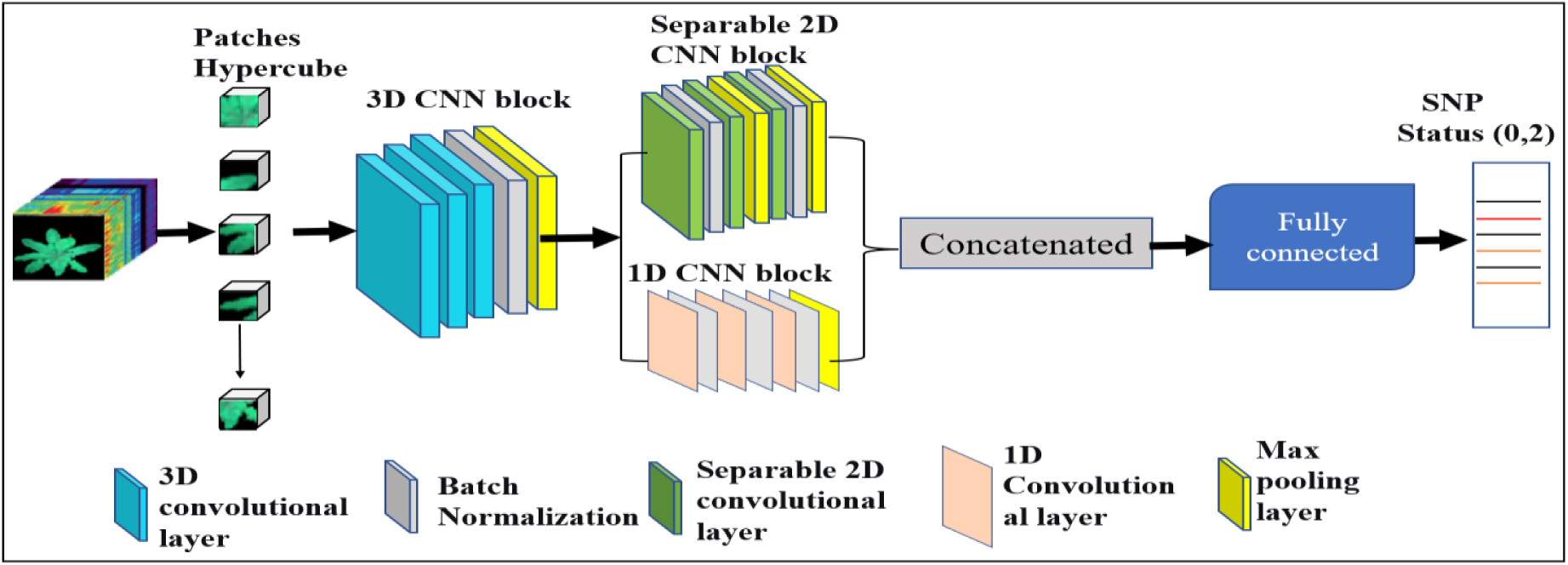
Architecture of the modified Hybrid-deep learning model for SNP identification from HSI data. A shared 3D CNN block extracts joint spatial-spectral features, which are then processed in parallel by a 2D CNN (spatial branch) and a 1D CNN (spectral branch). The outputs are concatenated and passed through a fully connected layer for binary classification.

The spectral branch (1D CNN) receives the 3D output and reshapes it into one-dimensional feature vectors, which are passed through three 1D convolutional layers (kernel sizes 3 and 5; filters = 8,16 and 32). This extracts detailed spectral features capturing biochemical and physiological properties, including pigment composition, anthocyanin accumulation, and stress-induced reflectance changes[27].

Batch normalization and dropout (0.3–0.5) were interspersed between the convolutions to stabilize training and reduce overfitting. A global average pooling layer was applied to each branch to compress features into compact representations. The outputs from the two parallel branches were concatenated and passed through a fully connected layer with a binary classification head. The model predicts homozygous reference (state 0) and homozygous alternate (state 2) genotypes. The full layer-by-layer architecture of the proposed hybrid model is provided in Supplementary Table S1.

#### 2.4.2 Developing the training dataset

To ensure unbiased model evaluation and prevent data leakage, the dataset was partitioned at the plot level. Each of the 194 genotypes was grown in two replicate plots, each containing 36 plants. For each plot, after excluding the border plants, **16** plants per plot were retained for further analysis. Plots were randomly split into 80% training (n = 310 plots) and 20% test (n = 78 plots). For each plant, five non-overlapping patches were randomly extracted, requiring at least 50% of plant pixels correspond to plant tissue based on an NDVI mask (**Supplementary Figure S3**). Each patch was assigned the SNP genotype label corresponding to its source plot, ensuring label consistency. This resulted in 124,000 training patches (310 plots × **16** plants × 5 patches × 5 time points) and 31,200 test patches (78 × 16 × 5 × 5). The same plot-level grouping was strictly preserved across all cross-validation folds, ensuring that patches from the same plot never co-occurred in training and validation sets. Within the training set, five-fold cross-validation was performed, corresponding to an 80:20 split between training and validation data in each fold (99,200 training patches and 24,800 validation patches per fold). To enhance spatial-spectral variability and improve generalization, 20% of training patches were augmented using geometric transformations (horizontal/vertical flipping and rotation) and additive noise.

#### 2.4.3 Hyperparameter optimization and training strategy

The hybrid CNN model was optimized using a hyperparameter tuning strategy to improve predictive performance and generalization. Key hyperparameters, including learning rate, batch size, number of filters, kernel size, and dropout rate, were optimized using five-fold grid search cross-validation. The Adam optimizer was used with learning rates ranging from 1×10^−5^ to 1×10^−2^. No weight decay was applied; regularization was achieved solely through dropout and early stopping. Batch sizes between 16 and 64 were evaluated to balance convergence stability and computational efficiency. Dropout rates in the range of 0.3 to 0.5 were applied for regularization. Early stopping was implemented with a patience of 10 epochs based on validation loss to prevent overfitting. Models were trained for up to 100 epochs, with learning rate scheduling applied to improve convergence. The grid search identified the optimal configuration as: batch size = 16, learning rate = 1×10^−4^, and dropout rates of 0.3–0.5 (with 0.5 applied to the final dense layer), selected based on the highest validation performance during cross-validation. The test set was strictly held out and used only for final performance evaluation to avoid optimistic bias.

#### 2.4.4 Model interpretation using Explainable AI

To interpret model’s predictions, two explainable AI (XAI) techniques were employed: SHapley Additive exPlanations (SHAP) and Gradient-weighted Class Activation Mapping (GRAD-CAM). SHAP analyzes the contribution of spectral features to model predictions based on cooperative game theory [28]. The analysis was performed on the fused latent feature representation from the concatenation of the spatial branch (2D CNN) and spectral branch (1D CNN) outputs prior to the final classification layer. The 3D CNN branch is progressively collapsed through the 2D and 1D branches before fusion; SHAP values corresponding to the 1D spectral branch were therefore isolated and aggregated to derive a global spectral importance profile, enabling direct attribution of prediction contributions to individual spectral bands. For a trained model *f*(*x*), where x ∈ ℝ^d^ represents the fused feature vector, the prediction is expressed as:

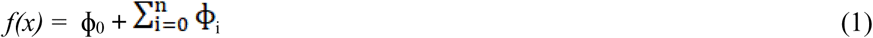

where ɸ_0_ is the expected model output over the background dataset and ϕi represents the contribution of the i-th feature to the prediction. SHAP values were computed using the DeepExplainer framework with 200 randomly selected training samples used as background data. To derive global spectral importance profile, SHAP values corresponding to the spectral branch were isolated and aggregated. Positive and negative SHAP values indicate features that increase or decrease the predicted probability of a given genotype class respectively.

Grad-CAM was applied to the final convolutional layer of the spatial branch to identify regions contributing to model predictions. For a target class c, with S_c_ been the score and A^k^, the K^th^ feature map, the importance weight is expressed as:

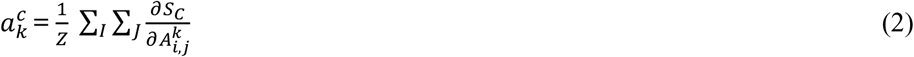

The resulting activation maps were upsampled to input resolution and overlaid on pseudo-RGB composites. Heatmaps were generated for both genotype classes and averaged across correctly classified samples to identify consistent spatial patterns, including leaf margins, venation structures, and localized pigmentation regions.

#### 2.4.5 Evaluation metrics for model performance

Model performance was evaluated using average accuracy (AA), F1 score, and receiver operating characteristic (ROC) curve analysis. AA provides a class-wise performance summary by computing the mean accuracy across classes. The F1-score accounts for class imbalance by combining precision and recall into a single metric. ROC analysis was critical because the workflow requires tuning the trade-off between sensitivity and specificity using different thresholds (e.g., high sensitivity for rare allele detection, high specificity for reliable genotyping). While AA provides an overview of per-class performance, the F1-score is more robust under class imbalance. ROC analysis complements both metrics by quantifying the model’s ability to distinguish between genotype classes across all thresholds. Together, these metrics provide a comprehensive evaluation of model robustness, sensitivity, and discriminative performance.

All models were trained on a high-performance computing server (HPC) at Utrecht University, equipped with NVIDIA GPUs (≥24 GB VRAM) and high-memory compute nodes. A single GPU was allocated per training job. Typical model training required approximately 1 hour, while full hyperparameter optimization was completed within several hours when executed in parallel. HSI preprocessing and model development were implemented in Python (v3.10) using TensorFlow with GPU acceleration via CUDA. Statistical analyses and SNP-level processing were conducted in R (version 4.2). This integrated Python-R pipeline provides a reproducible framework for HSI-based genotype classification, feature extraction, interpretation, and statistical validation.

## 3. RESULTS

This study investigates whether HSI captures biologically meaningful variation that enables the prediction of SNP genotypes in lettuce. To evaluate this, three distinct SNP categories were used. First, SNPs with known phenotypic impact — specifically, two anthocyanin-associated SNPs (ant.loc5 *and* ant.loc9), one leaf serration SNP (serr.loc5), and one pale pigmentation SNP (pale.loc4) — served as positive controls, as their effects on plant biochemistry and morphology are well established. Second, a null distribution was generated by randomly shuffling genotype labels to break any real genotype-phenotype association, providing a statistical baseline for chance performance. Third, 50 randomly selected SNPs with no known link to the measured phenotypes were used as a negative control set to assess model specificity and to potentially reveal uncharacterized associations.

Before addressing this SNP genotype prediction task, we first quantify spectral variability across genotypes to confirm that HSI data capture detectable differences among accessions. Subsequently, we evaluate the performance of the proposed hybrid deep learning model for SNP genotype prediction. (Note: this spectral analysis was performed prior to the SNP genotype prediction modeling)

### 3.1 Spectral Overview across Lettuce Cultivars

To characterize spectral variation among genotypes, spectral reflectance of twenty (20) randomly selected lettuce accessions were analyzed (for the whole 194 population, see Mehrem et al.[19]). All genotypes exhibited characteristic vegetation spectra, with strong absorption in the blue (approx. 400-500 nm) and red (approx. 660-680 nm) wavebands, a reflectance peak around 550 nm, and high reflectance in the near-infrared (NIR) regions. Although collectively the genotypes showed similar spectral characteristics, noticeable cultivar-specific variations were observed across the spectral range (Figure 5a).

**Figure 5.**
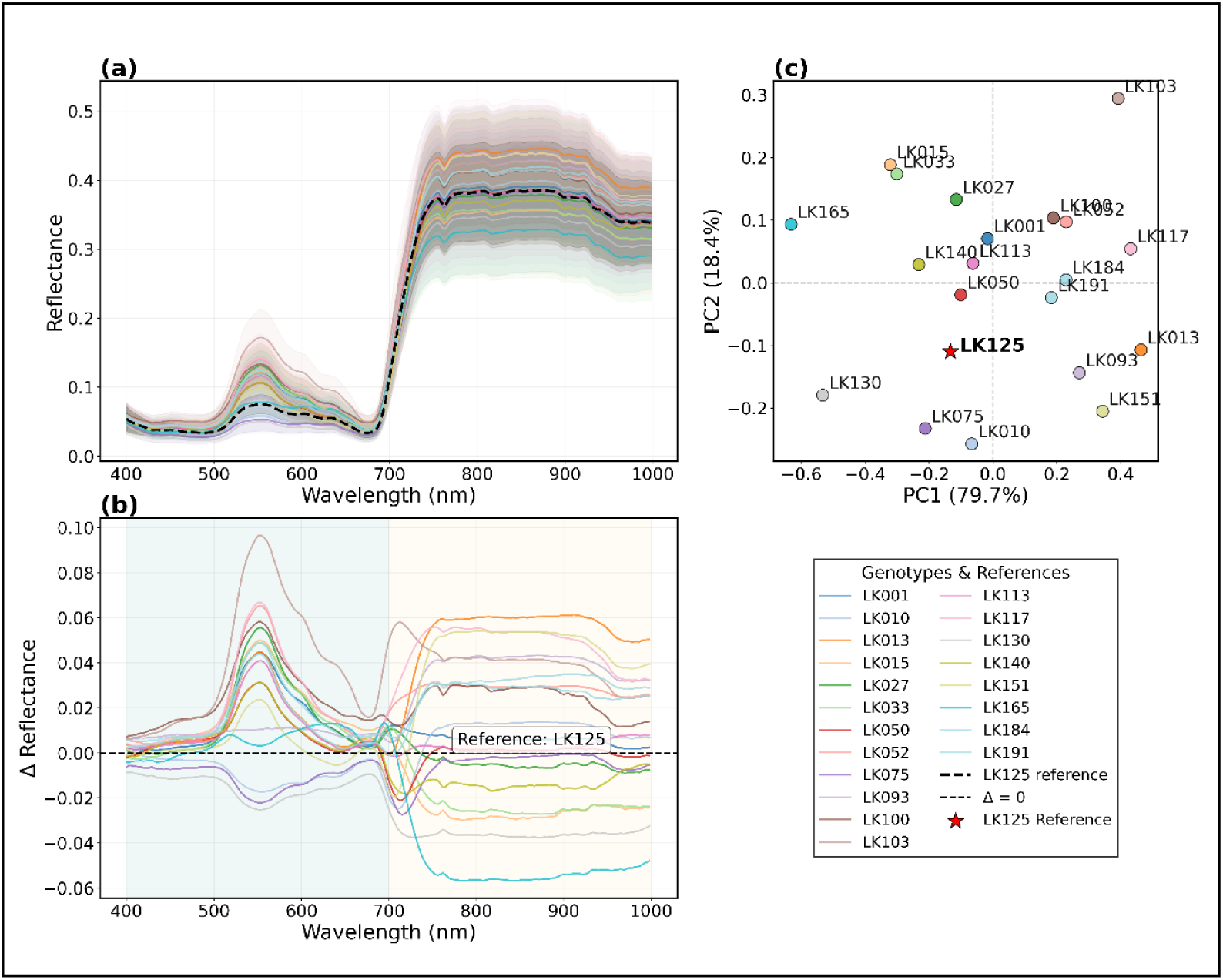
Spectral reflectance profiles and phenotypic variation in 20 selected Lettuce Genotypes (a) mean spectral curves with standard deviation for twenty randomly selected genotypes (b) difference spectra relative to reference genotype LK125; (c) PCA of spectral differences.

To quantify these differences, reflectance spectra of a reference genotype (LK125; Forellenschluss) were subtracted from those of other genotypes, producing difference spectra (Δ Reflectance) plotted against wavelength (Figure 5b). The cultivar LK125 was selected as the baseline reference because it represents the phenotypic extreme for anthocyanin accumulation in this lettuce population. Because high foliar concentrations of anthocyanin induce light absorption within the green-yellow spectral bands (∼520–580 nm), anchoring the analysis to this highly pigmented line maximizes the biological contrast against the greener accessions. This allows for a clear, biochemically anchored comparison that highlights the full gradient of pigment variation across the different cultivars. Several accessions exhibited clear deviations from the reference. For example, genotype LK027 (Ostinata) showed consistently lower Δ Reflectance across the visible spectrum 400-700 nm), with negative values ranging from –0.05 to –0.10, suggesting relatively higher pigment absorption. In contrast, genotypes LK015 (Capitan) and LK033 (Trgoviska) exhibited positive Δ Reflectance in the green-yellow bands (500-600 nm), peaking near 0.08 and 0.06 respectively. Conversely, genotype LK013 (Brenado) showed a mixed pattern: negative Δ Reflectance was observed in the blue region but positive Δ Reflectance showed in the NIR. LK050 (Diana) and LK075 (Mignonette) remained close to zero across most wavelengths, indicating spectra similar to the reference.

Principal component analysis (PCA) performed on the difference spectra showed that the spectral variability was dominated by the first two principal components, accounting for 97.7% of the total variance (PC1: 88.5%; PC2: 9.2%) (Figure 5C). PC1 was primarily associated with variation in the NIR region (above 700 nm), which could be influenced by leaf internal structure and water content. On the other hand, PC2 captured variation in the green spectral band (approximately 520-580 nm), mainly linked to carotenoid and chlorophyll ratio with the absorption peaking around 540 nm [29]. Plotting along PC1 and PC2, the twenty randomly selected genotypes did not form distinct clusters but instead showed a continuous spread. While genotype LK027 occupied the most negative extreme along PC2, LK015 and LK033 were positioned at the most positive region of PC2. Genotype LK125 (the reference) was located near the origin of the PC1-PC2 score plot, as expected. The remaining variance (2.3%) was distributed across higher-order principal components (PC3 and above), none of which individually exceeded 1.5% of explained variance.

### 3.2 Model Performance and Architecture Comparison

To evaluate the proposed hybrid model (3D-2D-1D CNN), its performance was compared with state-of-the-art architectures, including 3D-2D CNN, 2D CNN, and 1D CNN models, using the ant.loc5 SNP as the target label (Figure 6 and Table 1). The 3D-2D CNN achieved the highest predictive performance (accuracy: 0.938, F1-score: 0.915, AUROC: 0.925) but required longer training time (3150 s) and exhibited considerable variability between runs (**Figure 6a**). The proposed hybrid model delivered competitive performance (accuracy: 0.922, F1-score: 0.891, AUROC: 0.906) while reducing training time by 33.7% (2088 s) and demonstrating superior stability, with the narrowest interquartile range and lowest standard deviation across repeated runs. The 2D and 1D CNN models were computationally efficient (<670 s) but showed substantially lower predictive performance and higher variability, making them less suitable for robust phenotyping applications.

**Figure 6.**
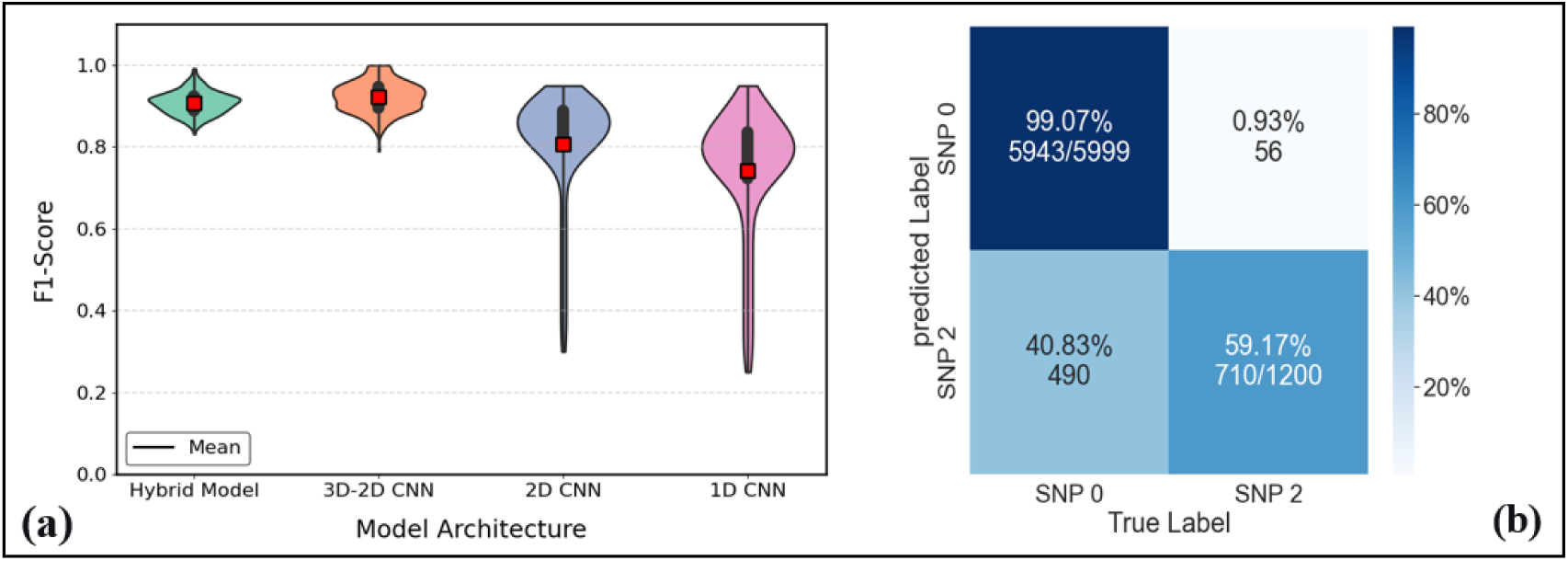
Performance comparison of CNN architectures for SNP genotype prediction using ant.loc5 SNP; (a) violin plots showing the distribution of F1-scores across repeated runs (red squares indicate the mean score); (b) confusion matrix of the hybrid CNN model illustrating class-wise prediction outcomes under high class imbalance.

**Table 1.** Comparative performance of hybrid CNN model and state-of-the-arts CNN using ant.loc5. Metrics include AA, F1-score, AUROC, and Training Time (in seconds). Values represent mean ± standard deviation.

| Model | AA | F1-score | AUROC | Training Time (s) |
| --- | --- | --- | --- | --- |
| Hybrid CNN model | 0.922±0.018 | 0.891±0.015 | 0.906±0.026 | 2088.06±75.44 |
| 3D-2D CNN | 0.938±0.047 | 0.915±0.094 | 0.925±0.025 | 3150.01±29.98 |
| 2D CNN | 0.891±0.172 | 0.869±0.227 | 0.817±0.200 | 662.96±25.095 |
| 1D CNN | 0.853±0.244 | 0.804±0.235 | 0.813±0.280 | 605.06±28.472 |

To understand the error profile of the hybrid model, confusion matrix analysis (**Figure 6b**) revealed that the majority class (SNP 0) was correctly classified with an error rate of 0.4%, while approximately 41.83% of the minority class was misclassified. This disparity reflects the inherent class imbalance in the ant.loc5 dataset, where SNP 0 (homozygous reference) constitutes the majority class. Despite this, the high AUROC of 0.906 ± 0.026 indicates strong overall discriminative ability, as AUROC is insensitive to class imbalance unlike accuracy-based metrics In contrast, the 2D and 1D CNNs (**Supplementary Figure S4**) exhibited higher misclassification rates, particularly for the minority class, where correct identification dropped below 50% in some cross-validation folds.

### 3.3 Performance analysis of the SNPs modeling

Next, we evaluated model performance using additional SNPs with known phenotypic impact and a set of 50 randomly selected SNPs (see Figure 7). By comparing model performance against a null distribution (obtained via random label permutation), we assessed whether high performance reflects hyperspectral phenotype–genotype associations. This analysis establishes the reliability and biological relevance of the proposed framework.

**Figure 7.**
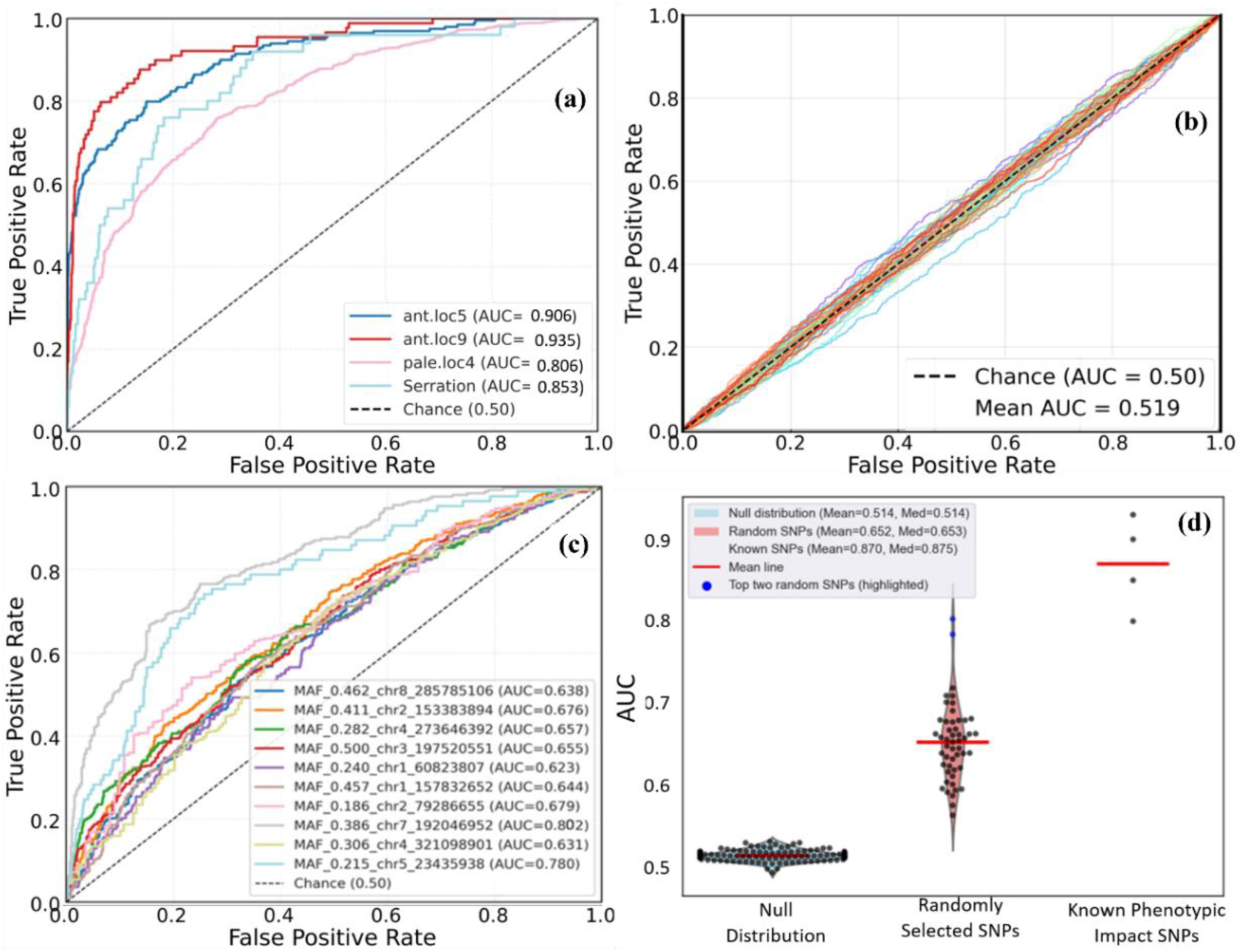
Comparative analysis of classification performance: (a) ROC curves for SNPs with known phenotypic impact; (b) null distribution under random labeling; (c) representative ROC curves for randomly selected SNPs; (d) violin plots of AUC distributions across SNP categories. (Supplementary Figure S5 shows the remaining randomly selected SNPs ROCs).

#### 3.3.1 SNPs with Known Phenotypic Impact

Evaluation of the proposed hybrid model on SNPs with known phenotypic impact provides a benchmark for detecting genotype-driven spatial-spectral variation. Models trained on these SNPs demonstrated strong predictive performance (Figure 7a and Table 2). Model with **ant.loc9** achieved the highest accuracy (0.951 ± 0.011), followed by ant.loc5 (0.922 ± 0.018), serr.loc5 (0.860 ± 0.065), and pale.loc4 (0.833 ± 0.052). All accuracies substantially exceeded the null expectation (mean ∼0.51). The models also demonstrated balanced precision-recall trade-off, as reflected in high F1-scores, with ant.loc9 achieving the highest (0.914 ± 0.070). This was followed by ant.loc5 (0.891 ± 0.015) with pale.loc4 (0.813 ± 0.046) achieving good but slightly lower performance. The serr.loc5 SNP yielded an F1-score of 0.830 ± 0.352; the larger standard deviation suggests higher sensitivity to class imbalance or sample variation.

**Table 2.** Model performance for known phenotypic SNPs and four randomly selected SNPs (two high and low performing SNPs). Metrics include Average Accuracy(AA), F1-score, AUROC. Values represent mean ± standard deviation.

| Group Name | SNP | AA | F1-score | AUROC |
| --- | --- | --- | --- | --- |
| Known Phenotypic | ant.loc5 | 0.922±0.018 | 0.891±0.015 | 0.906±0.026 |
| Impact SNPs | ant.loc9 | 0.951±0.011 | 0.914±0.070 | 0.935±0.142 |
| | pale.loc4 | 0.833±0.052 | 0.813±0.046 | 0.806±0.197 |
| | serr.loc5 | 0.860±0.065 | 0.830±0.352 | 0.853±0.024 |
| Randomly Selected | MAF_0.386_chr7_192046952 | 0.822±0.026 | 0.785±0.033 | 0.802±0.042 |
| SNPs (high-performing) | MAF_0.215_chr5_23435938 | 0.791±0.063 | 0.783±0.024 | 0.780±0.031 |
| Randomly Selected | MAF_0.381_chr4_268104397 | 0.5811±0.05 | 0.5671±0.039 | 0.5631±0.045 |
| SNPs (low-performing) | MAF_0.302_chr9_123738720 | 0.595±0.066 | 0.560±0.025 | 0.575±0.018 |

AUROC scores further confirmed strong discriminative power, with ant.loc9 achieving the highest (0.935±0.142), followed by ant.loc5 (0.906±0.026), serr.loc5 (0.853±0.024), and pale.loc4 (0.806±0.197). The relatively higher standard deviation for for ant.loc9 and pale.loc4 suggests some variability across cross-validation folds, possibly due to class imbalance or population structure. Nevertheless, all AUROC values were significantly above the 95th percentile of the null distribution, which suggest true genotype-phenotype association.

The prediction probability distribution (Figure 7d) showed that all four points (SNPs with known phenotypic SNPs) formed a distinct cluster at the upper AUC scale and well separated from the null distribution (centered around 0.514). The clear elevation of SNPs with known phenotypic effect indicates consistent and robust model performance.

#### 3.3.2 Null Distribution (Random Labeling)

To determine whether model performance arises from meaningful genotype-phenotype associations rather than overfitting or high-dimensional artifacts, a null distribution was generated by randomizing genotype labels while preserving class balance and sample size. This strategy removes the true genotype–phenotype association, providing a statistical baseline for assessing non-spurious learning. Under random labeling, the model exhibited low discriminative power. As shown in Figure 7b, the ROC curve closely followed the diagonal chance line (with true positive rate ≈ false positive rate) with a mean AUROC ≈ 0.50. The AUC distribution (Figure 7d) was approximately normal and centered around 0.50. This reflects the absence of learnable patterns, confirming that without true genotype-phenotype association, the model cannot extract consistent signals from the HSI data.

The 95^th^ percentile of the null distribution (AUC= 0.514) was used as a statistical threshold (α < 0.05) to assess significance. Any true SNP whose model achieves an AUROC above this threshold suggest non-random genotype-phenotype association. This threshold provides a robust baseline for evaluating candidate SNPs, including those with previously unknown phenotypic effects. The specificity of the null distribution approach is critical as it accounts for potential overfitting, class imbalance, and the high dimensionality of HSI data. The demonstration that random labels yield near-chance performance validate that the model’s strong performance on known phenotypic SNPs or on the randomly selected SNPs is driven by true biological signals rather than noise or spurious correlations.

#### 3.3.3 Randomly Selected Putatively Neutral SNPs

In this section, we evaluated model performance on randomly selected SNPs with no known phenotypic association to assess model specificity and explore potential unexpected genotype–phenotype signals. Overall, most randomly selected SNPs exhibited lower and more inconsistent performance compared to SNPs with known phenotypic impact (Figure 7c; Supplementary Table S2). The mean accuracy across randomly selected SNPs is 0.607 ± 0.072, which is above the null baseline. AUROC scores range from 0.500 to 0.790, with most randomly selected SNPs falling below 0.700. The AUROC distribution (Figure 7d) was bimodal and skewed, with a mean of 0.607 and median of 0.593. This indicates two distinct groups: (i) a dominant group of low-performing SNPs (AUROC < 0.62), consistent with weak or absent phenotypic signal, and (ii) a smaller subset of high-performing SNPs (AUROC > 0.72). Hence, the randomly selected SNPs set ranges from weak association to moderate association with HSI phenotypes.

Among the randomly selected SNPs, the top two SNPs (MAF_0.386_chr7_192046952 and MAF_0.215_chr5_23435938) achieved AUROC scores between 0.780 and 0.802, which were near the lower end of the performance range for SNPs with known phenotypic impact (0.823-0.930). These SNPs were selected for further analysis to investigate whether their predictive signal reflects residual confounding or potentially uncharacterized genotype-phenotype relationships.

Collectively, the predominance of the low-performing SNPs shows that most randomly selected SNPs have little measurable phenotypic effects in the data. This further confirms that the model’s strong predictive performance on SNPs with known phenotypic impact is driven by biologically meaningful patterns in the HSI data, not by noise or overfitting.

### 3.4 Interpretation of the Genotype-Phenotype Associations

To determine whether model predictions are driven by meaningful spatial-spectral features, we employed two explainable AI techniques, GRAD-CAM and SHAP analysis [30].

#### 3.4.1 SHAP Analysis for Spectral Importance

The important wavelengths for ant.loc5 were embedded in the visible regions (508 nm - 538 nm), red-edge (704.53 nm) and NIR (701.81 and 704.53 nm) (***Figure 8a***). Most of the contributions were positive, indicating reflectance in those bands is associated with anthocyanin-accumulation. This aligns with known anthocyanin absorption in the green-yellow region. The wavelengths identified by SHAP as most influential for anthocyanin-related SNPs (508–538 nm, 704 nm) align with the spectral variation captured by PC2 in Figure 5c, confirming that pigment-related reflectance differences drive both global spectral variability and model predictions. Similarly, ant.loc9 was influenced by wavelengths in green-yellow region (516.29-566.94 nm) and the NIR region (902 nm - 936.38 nm) (Supplementary Figure S6a). However, some of the NIR wavelengths showed negative SHAP values (e.g. 925.19 nm), indicating a more complex relationship of SNPs influencing biochemical processes. Pale.loc4 showed a bidirectional pattern moving from the visible to the NIR wavebands with 516.29 and 566.94 dominating in the visible regions while 925.19 nm and 936 nm were strong NIR contributors (Supplementary Figure S6b). The serr.loc5 SNP showed strong positive SHAP values localized within the visible region (540 - 550 nm) and extended into the red-edge (696.37 nm) and NIR (734.52 nm) (Supplementary Figure S6c). The high performing randomly selected SNP (MAF_0.386_chr7_192046952) displayed a distinct SHAP profile dominated by the NIR region (701.81 - 704.53 nm). The positive contributions in the NIR region suggest that this SNP may influence leaf internal structure and biochemical processes, which requires further investigations. In contrast, the low predictive performance displayed diffuse SHAP distributions with no dominant wavelengths and mixed feature gradients. This indicates weak and non-specific spectral feature utilization.

**Figure 8.**
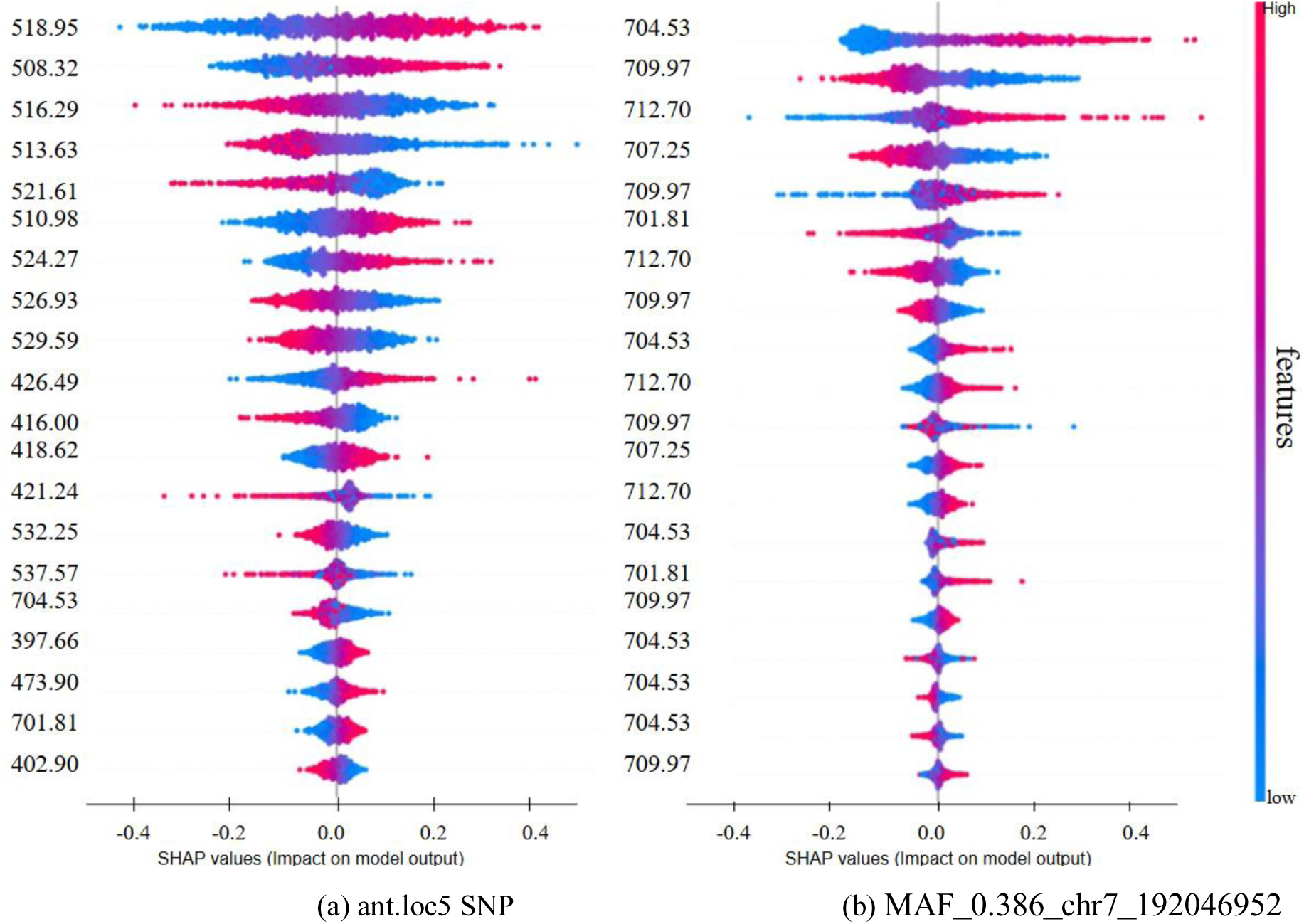
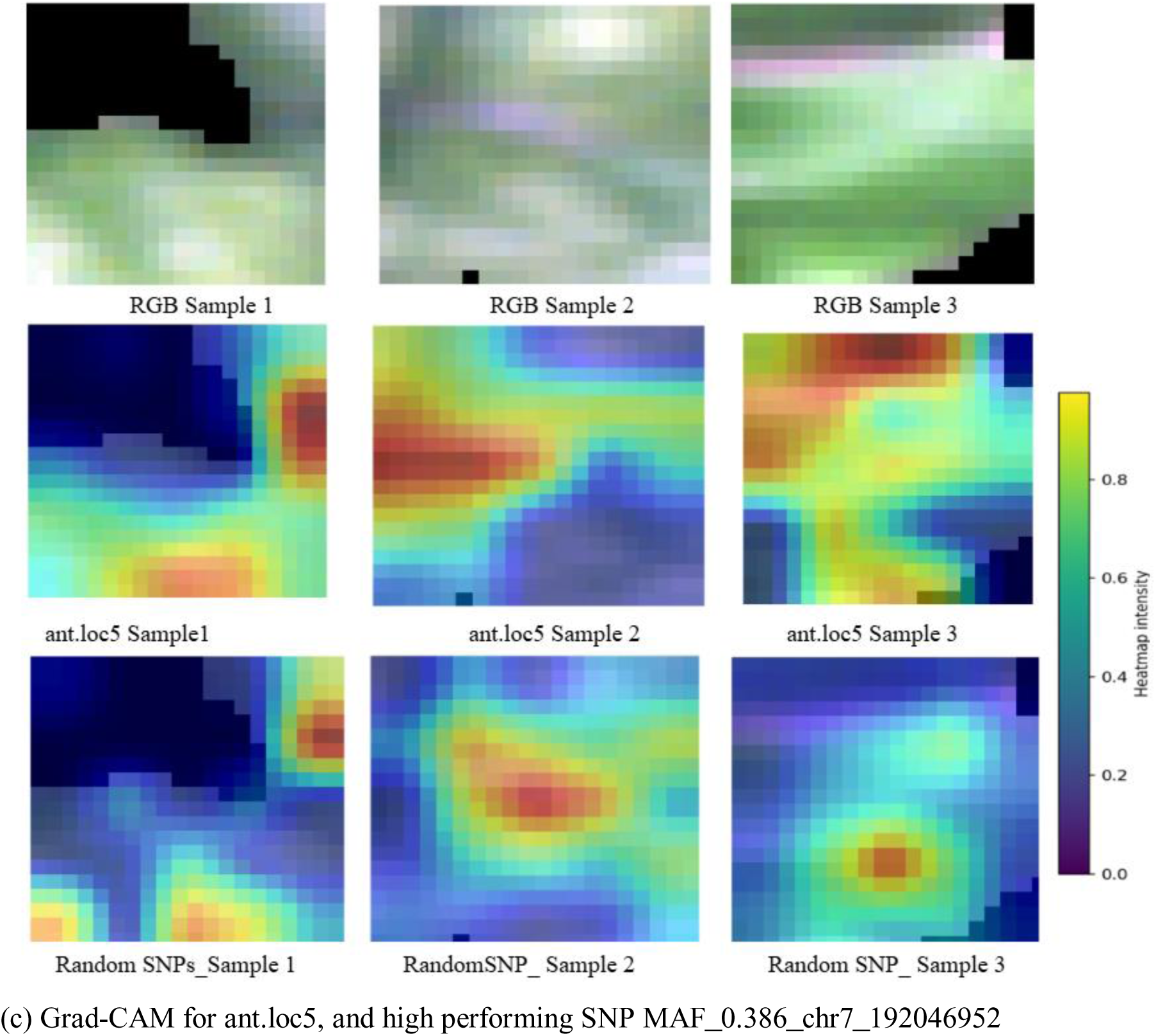
Explainable AI interpretation of model predictions. (a) SHAP-based spectral importance for SNPs with known phenotypic impact (ant.loc5); (b) SHAP-based spectral importance for the high-performing randomly selected SNP (MAF_0.386_chr7_192046952). Positive SHAP values indicate that higher reflectance at those wavelengths increases the prediction probability of the alternate genotype (SNP 2), while negative values support the reference genotype (SNP 0). (c) GRAD-CAM spatial activation maps for models with varying performance: pseudo-RGB images (top row), ant.loc5 (middle row), and the randomly selected SNP MAF_0.386_chr7_192046952 (bottom row).

The SHAP profiles for all the high performing SNPs (including SNPs with known phenotypic impact) were structured and reproducible across samples, confirming that the model learned biologically meaningful spectral signatures. In contrast, low-performing random SNPs (Figure S6d) produced flat, noisy SHAP distributions with inconsistent wavelength dominance, confirming the weak signals or absence of a consistent genotype-phenotype association.

#### 3.4.2. GRAD-CAM for Spatial Attribution

GRAD-CAM analysis was applied to interpret spatial feature contributions learned by the model. The analysis revealed distinct spatial activation patterns across SNP categories (Figure 8b; Supplementary Figure S7). SNPs with known phenotypic impact exhibited structured and biologically relevant spatial attention with activations concentrated within plant tissues. This suggests that predictions were driven by biologically relevant patterns rather than background artifacts. The serr.loc5 SNP exhibited activation localized along leaf margins, consistent with morphological variation in leaf edges. The low-performing randomly selected SNPs showed cluttered and unstructured patterns with near-zero activation across the image, suggesting minimal spatial contributions to the model’s prediction. However, the high-performing randomly selected SNPs exhibited more structured activation within plant regions, aligning with their corresponding spectral importance patterns observed in SHAP analysis. Together, these results demonstrate that the hybrid model captures complementary spectral and spatial features. Strong predictive performance is associated with consistent wavelength-specific importance and structured spatial activation, whereas weak performance corresponds to diffuse and non-specific feature attribution.

#### 3.4.3 Validation across extended SNP Set

To further validate these findings, we examined the trained model’s predictions on heterozygous samples (genotype 1). This analysis empirically justified our binary classification approach and provided biological insight into allelic dominance, as the model’s continuous probability outputs reflected genetic dosage effects. Post-hoc examination revealed distinct locus-specific patterns: ant.loc9 heterozygotes clustered near the homozygous reference class, while pale.loc4 heterozygotes displayed a broad distribution spanning both homozygous classes. No heterozygotes were present for ant.loc5 *or* serr.loc5. Validation across varying sample sizes and MAF thresholds (Supplementary Figure S8) confirmed consistent performance, indicating that high-performing SNPs reflect genuine genotype-phenotype associations rather than sampling artifacts.

## 4. DISCUSSION

This study presents a deep learning framework linking genetic variation to high-dimensional phenotypic signatures by inferring SNP genotypes directly from hyperspectral imaging (HSI) data. Prior to model development, spectral profiles were analyzed to assess genotype-dependent variation. Although average spectral profiles across genotypes appeared broadly similar, subtle differences in spectral shape and intensity across wavelengths enabled discrimination among genotypes. These variations are consistent with established relationships between reflectance characteristics and plant physiology. For instance, variation in the red-edge and near-infrared regions is associated with internal leaf structure and light scattering within the mesophyll, while visible wavelengths are primarily linked to pigment absorption[29]. Furthermore, reflectance variation is influenced by canopy architecture, which affects the near-infrared plateau through changes in scattering dynamics [31][32].

These subtle yet informative variations highlight the need for modeling approaches capable of capturing complex spatial-spectral relationships. From a methodological perspective, the hybrid model demonstrated that jointly modeling spatial and spectral information improves predictive performance. Compared to state-of-the-art models, the hybrid approach reduced variance and improved generalization, suggesting that genotype–phenotype relationships are inherently complex and nonlinear. Furthermore, the reduced computational cost of the hybrid model relative to 3D convolutional approaches makes it more suitable for scalable phenotyping pipelines[33].

By leveraging both spectral and spatial information, the proposed framework enables the prediction of SNP genotypes from HSI-derived phenotypes, providing a complementary perspective to conventional genotype–phenotype mapping. The high predictive performance observed for SNPs with known phenotypic effects, particularly those associated with anthocyanin-related loci (e.g., *ant.loc5* and *ant.loc9*), supports the hypothesis that functional variants affect biochemical processes, which in turn generate consistent and learnable spectral patterns in HSI data. In contrast, neutral variants do not produce distinguishable signatures.

The ability of the model to accurately predict specific SNP genotypes from HSI data highlights the sensitivity of hyperspectral measurements to underlying biochemical changes. Anthocyanin accumulation, for example, is known to influence reflectance properties in the visible and near-infrared regions, particularly within wavelengths associated with pigment absorption[34]. The high predictive performance for SNPs with known phenotypic effects therefore provides biological validation of the framework, suggesting that the model captures physiologically meaningful variation rather than spurious correlations. Similarly, the detection of SNPs associated with leaf structural traits indicates that HSI can encode subtle morphological and anatomical differences through spatial-spectral patterns.

In addition to predictive modeling, this study introduces a statistical strategy for evaluating genotype-phenotype associations through null distribution estimation via label randomization. This approach enables differentiation between biologically meaningful signals and random patterns, providing a baseline for interpreting model performance. Model performance exceeding this baseline is unlikely to arise from overfitting or random correlations. SNPs with known phenotypic effects consistently exceeded this threshold, supporting the validity of this method. In contrast, most randomly selected SNPs exhibited performance close to the null baseline, indicating little or no detectable signal in HSI data for these SNPs.

In contrast, a small subset of randomly selected SNPs showed relatively high predictive performance (AUROC > 0.70), suggesting the presence of measurable signals. To rule out residual linkage disequilibrium as an explanation for the high predictive performance of these two SNPs, we computed LD (r²) between them and the four known phenotypic SNPs. All pairwise r² values were well below our exclusion threshold of 0.3 (maximum r² = 0.131), confirming that the observed signals are unlikely to be driven by linkage to the known functional loci. However, these observations should be interpreted with caution. Despite efforts to minimize LD with known loci, residual LD with unknown functional variants cannot be ruled out. Alternatively, these SNPs may capture subtle phenotypic effects not detected by conventional GWAS approaches, highlighting the sensitivity of HSI to underlying physiological variation. In this case, model predictive performance may serve as a proxy for biological relevance, although further validation is required.

A key contribution of this work lies in the integration of explainable artificial intelligence (XAI) techniques to interpret model predictions. SHAP and Grad-CAM analyses provide a bridge between data-driven modeling and biological interpretation, enabling the identification of candidate spectral features linked to specific physiological processes. The identification of anthocyanin-related SNPs aligns with established spectral signatures of anthocyanin and chlorophyll-related traits [35][6]. Similarly, the mapping of the pale.loc4 SNPs to the chlorophyll absorption bands (400-650 nm) is consistent with its role in green pigmentation[36]. Grad-CAM further revealed spatially coherent activation patterns localized within plant tissues, indicating that predictions were driven by biologically relevant regions rather than background artifacts. In contrast, low-performing SNPs exhibited diffuse and unstructured SHAP and Grad-CAM patterns, consistent with the absence of meaningful genotype–phenotype signals. Notably, high-performing randomly selected SNPs displayed more structured spectral and spatial patterns, suggesting that the model captured consistent, non-random variation. However, whether these patterns reflect true biological effects or residual confounding requires further investigation. Consequently, this framework is best suited as a targeted screening tool for pre-selected subsets of variants or SNPs within functionally annotated genomic regions. This strategy reduces computational burden while preserving biologically relevant candidates. Furthermore, the ability to infer genotype from phenotype opens possibilities for reducing reliance on traditional genotyping in specific contexts, particularly where rapid and non-destructive screening is required.

Beyond the XAI analysis the trained model was also used to examine the heterozygous exclusion decision post-hoc. The predicted probability distributions of heterozygous individuals (Supplementary Figure S8) confirmed that for ant.loc9, predicted probabilities clustered near zero– difficult to distinguish from the homozygous reference class. This suggests that a single alternate allele at this locus is insufficient to produce a detectable spectral shift. In contrast pale.loc4 exhibited a broad distribution spanning both homozygous reference (0) and alternate (2) classes. This indicates incomplete dominance (additive effect) in chlorophyll regulation. For ant.loc5 and the serr.loc5 SNPs, no heterozygous individuals were present in the dataset, precluding this analysis. Together, these observations support our binary classification formulation: heterozygous samples either collapse into one homozygous class or show an ambiguous intermediate signal that would degrade classification performance. Excluding them from training thus avoids degraded classification performance and ambiguity at the decision boundary. The distinct distributional patterns observed across loci also suggest that the model’s predicted probabilities could serve as a proxy for inferring allelic dominance relationships — this is worth exploring in future works.

Despite these strengths, some limitations should be acknowledged. Applying this approach to genome-wide SNP screening is computationally expensive. A conservative back-of-the-envelope estimate suggests that evaluating all 2.3 million SNPs would require millions of GPU-hours (It takes over 2100 seconds to train and evaluate on SNP), rendering it infeasible. Secondly, the framework is dependent on high-quality, well-aligned HSI and genotype data, and its performance may be affected by noise, environmental variability, or inconsistencies in data acquisition. Future work could address these limitations by incorporating multi-environment datasets, integrating temporal dynamics, and combining dynamic modeling approaches with data-driven approaches. Additionally, class imbalance across some SNP genotype classes (e.g. ant.loc5) represents a practical limitation for applications requiring balanced sensitivity across genotype classes. Future work could address this through cost-sensitive learning, oversampling strategies such as SMOTE, or threshold adjustment based on the predicted probability distributions.

In conclusion, this study demonstrates that HSI data, when combined with deep learning and explainable AI, *c*an be used to detect genetic variation signatures in phenotypic data*, enabling* hypothesis-driven screening of genotype–phenotype associations. By establishing a direct and interpretable link between genotype and high-dimensional spatial-spectral phenotypes, the proposed framework advances the state of the art in plant phenomics and provides a foundation for future research in data-driven crop improvement.

## 5. CONCLUSION

This study demonstrates that hyperspectral imaging (HSI), combined with deep learning, can be used to **detect** genotype--phenotype relationships *by* predicting genotypes at SNP loci from hyperspectral phenotypes *-* a reverse paradigm compared to conventional approaches. The hybrid CNN framework achieved high predictive accuracy for SNPs with known phenotypic effects, while a statistical null distribution confirmed that this performance reflected genuine biological signals rather than noise. Explainable AI further linked model predictions to interpretable spectral and spatial features, bridging the gap between data-driven modeling and plant physiology.

Beyond demonstrating feasibility, this framework has practical implications for crop breeding and genetic discovery. By enabling non-destructive prediction of SNP genotypes from HSI data, DeepPheno offers a pathway for rapid, high-throughput screening of breeding populations without the need for extensive genotyping. The ability to identify genotype–phenotype associations from imaging data also positions this approach as a hypothesis-generating tool for discovering novel traits and candidate genes. In the future, integrating this framework with temporal HSI data, multi-environment trials, and genomic prediction models could further accelerate marker-assisted selection and precision breeding. With validation and scaling, DeepPheno has the potential to become an AI tool in plant breeding pipelines, complementing genomic selection by providing cost-effective, field-deployable phenotypic profiling.

## Authorship contributions

FGO and SA conceived the study, SLM, GvdA and BLS collected the data and phenotypes, FGO, BLS and SLM SLM processed and analyzed the data. FGO performed the AI work. FGO wrote the manuscript with input from all co-authors. All authors read and approved the final manuscript.

## Data Availability

The code, scripts, and supplementary material supporting the findings of this study have been deposited in the GitHub repository at https://github.com/frankgyan/Utrecht-University--HSI. The raw source datasets analyzed during this study are proprietary commercial property shared under a non-disclosure agreement by a consortium of nine seed companies; therefore, they cannot be made publicly available due to legal and confidentiality restrictions. To facilitate reproducibility and code verification, a downscaled, de-identified sample dataset has been provided within the repository to demonstrate script execution and pipeline workflow. Access to the full underlying commercial dataset must be requested directly from the participating seed companies and is subject to consortium approval.

## Declaration of competing interests

The authors declare that they have no known competing interests (financial nor personal relationships) that could have appeared to influence the work reported in this paper.

## Acknowledgements

This research is part of the LettuceKnow project (project number 1.2 of the research Perspective Program P19-17). The project is partly sponsored by the Dutch Research Council (NWO), the associated breeding companies: BASF, Bejo Zaden B.V., Limagrain, Enza Zaden Research & Development B.V., Rijk Zwaan Breeding B.V., Syngenta Seeds B.V., and Takii and Company Ltd., and the Foundation for Food and Agriculture Research (FFAR).

## Notes

### Competing Interest Statement

The authors have declared no competing interest.

