## Supplementary Information for "DeepPheno: A Deep Learning Framework for Linking Hyperspectral Imaging and SNP Genotypes in Lettuce"

### SUPPLEMENTARY MATERIALS

#### Figure S1- S8

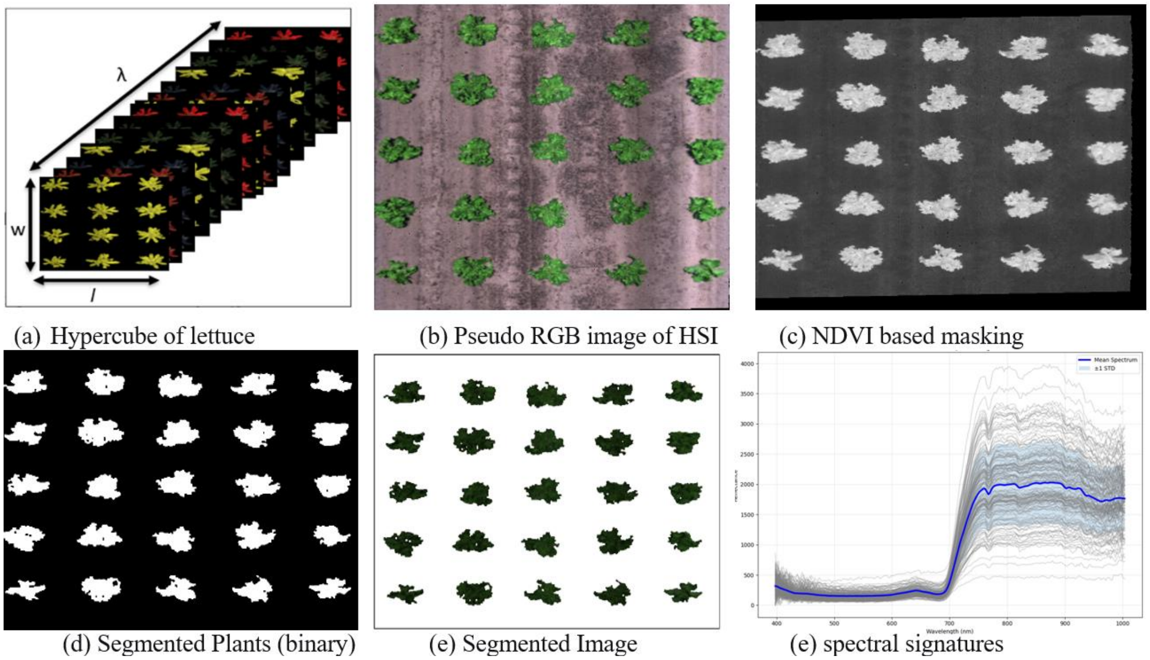


**Figure S1.** Hyperspectral image segmentation steps

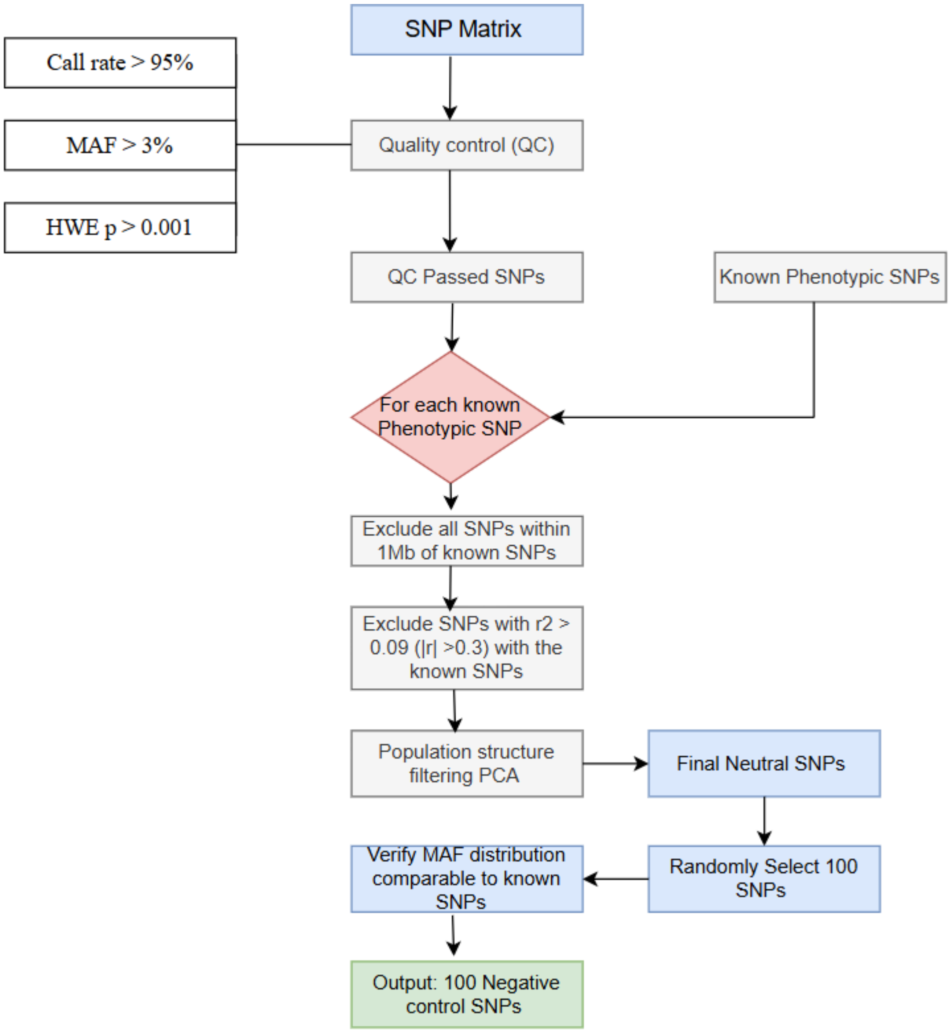


**Figure S2.** Pipeline to extract randomly selected SNPs from the genome matrix

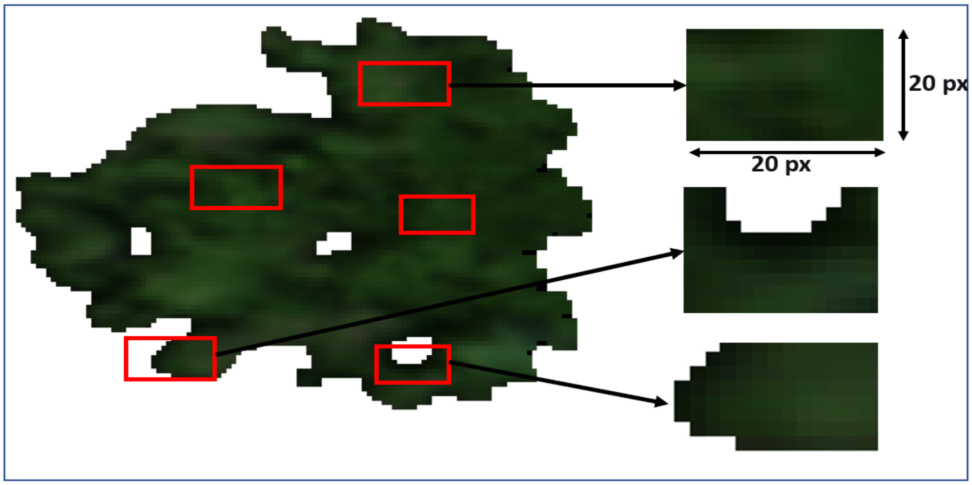


**Figure S3.** Patches Extraction from Hyperspectral Imaging

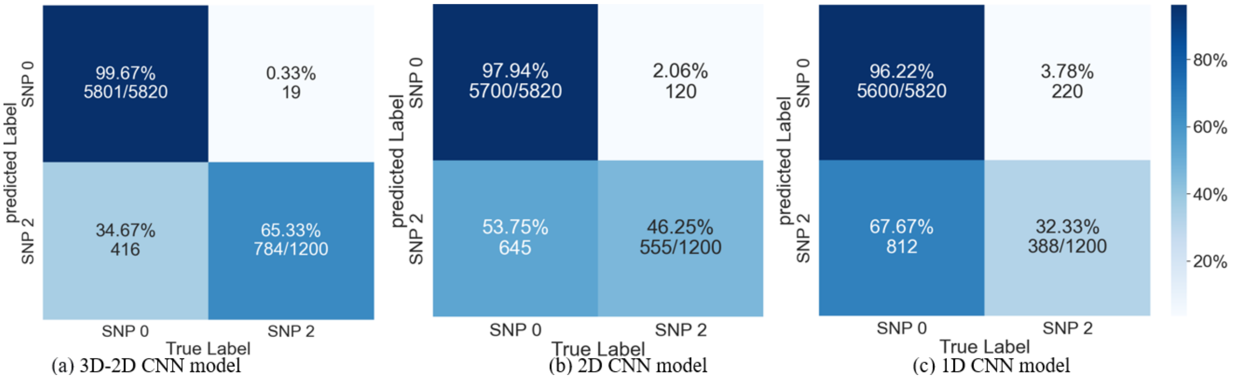


(a) 3D-2D CNN model (b) 2D CNN model (c) 1D CNN model

**Figure S4:** Confusion matrices for the SNPs with known phenotypic impact

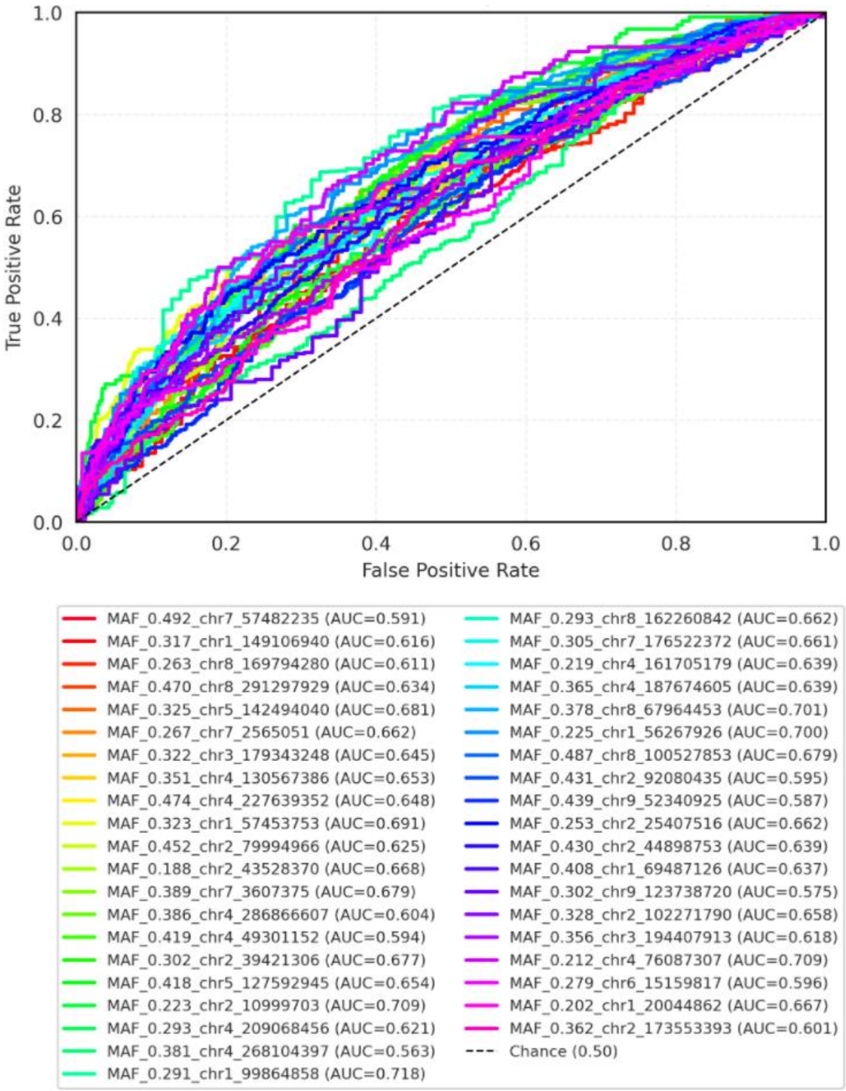


**Figure S5.** Representative ROC curves for randomly selected SNPs for the other 40 selected SNPs

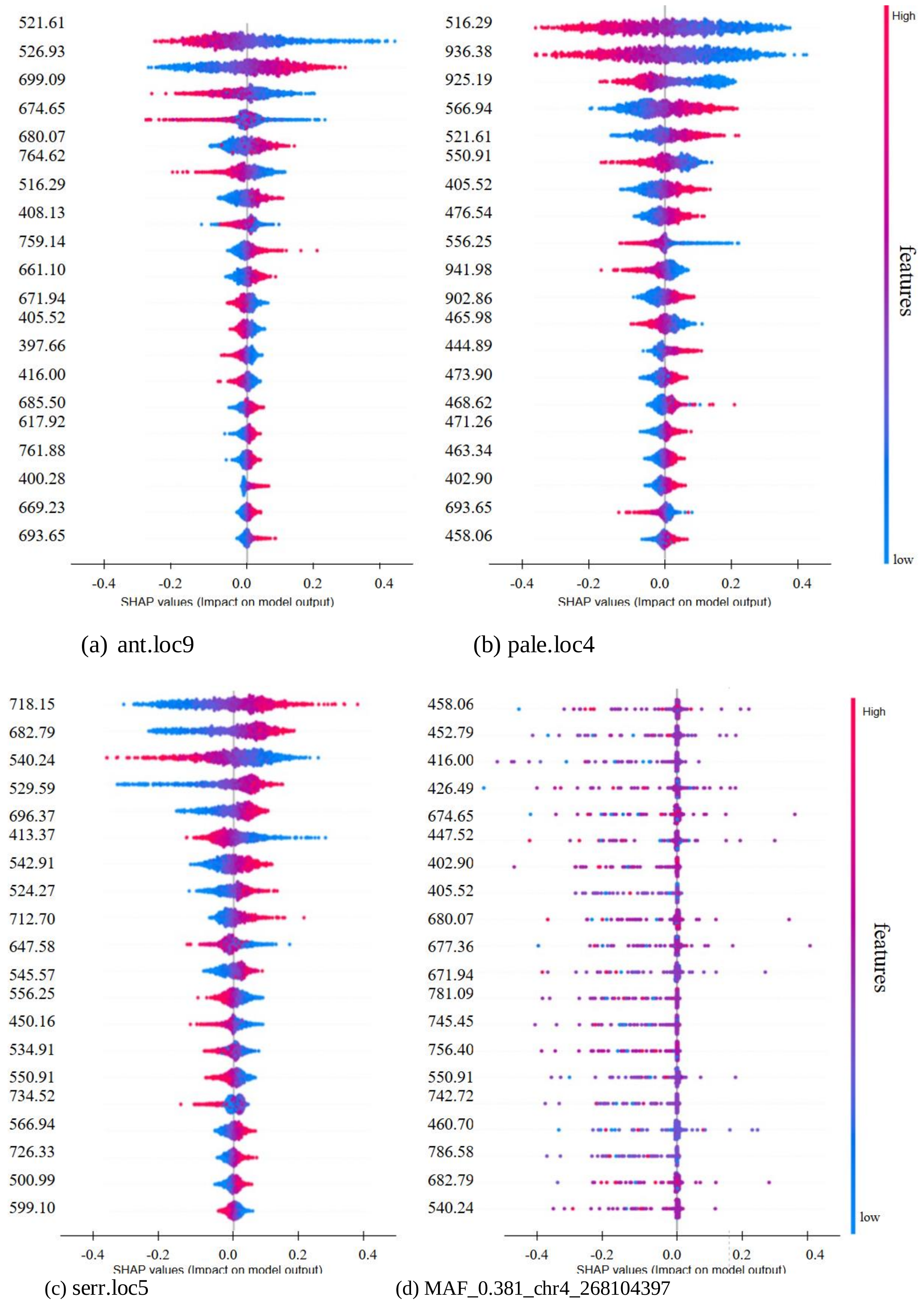


**Figure S6.** Visualization of model performance based SHAP plots of the SNPs with known phenotypic impact models

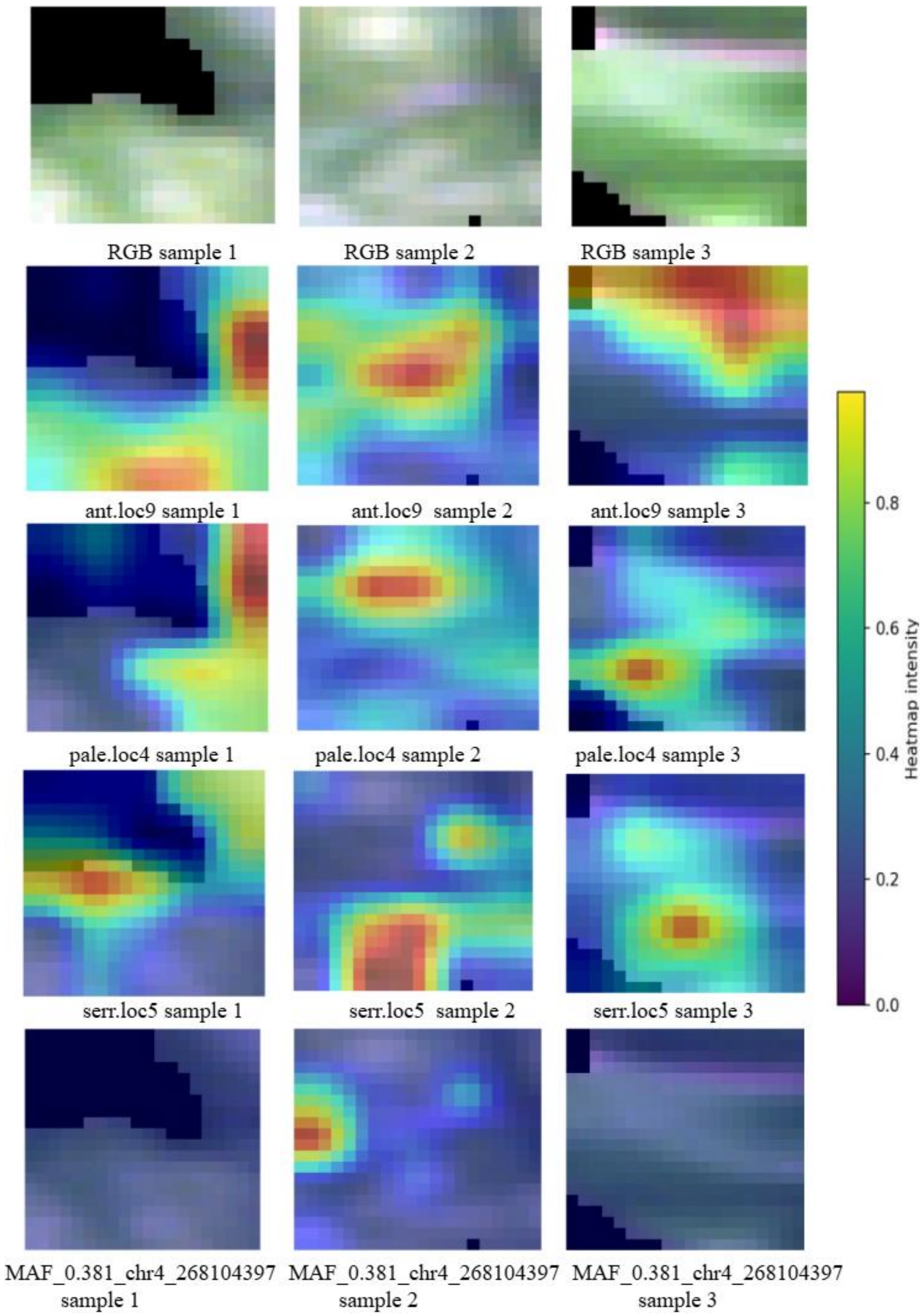


**Figure S7.** Visualization of model performance based GRAD-CAM of randomly selected patches of the SNPS with known phenotypic impact models (first column are the grey scale images, second third and fourth columns are GRAD-CAM for of ant.loc9, pale.loc4 and *serr.loc5* models respectively.

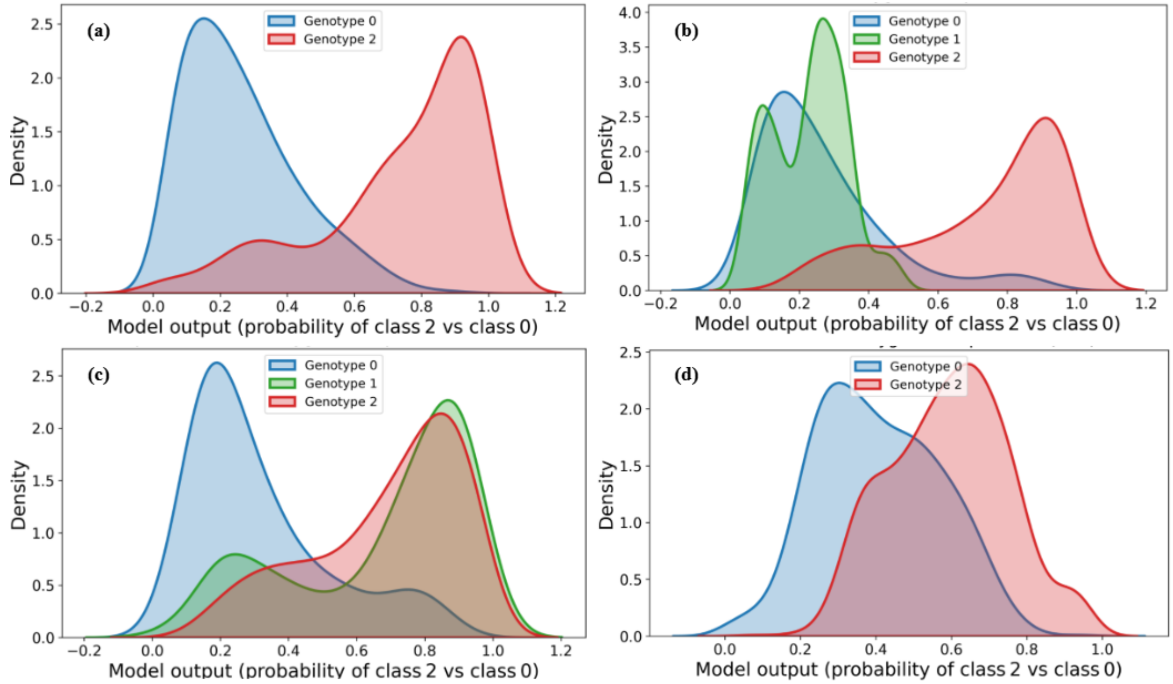


**Figure S8.** Heterozygote interpretation for four functional SNPs**.** Each panel shows the kernel density estimate of the model output for genotypes 0, 1, and 2 (blue, green, red). The model was trained only on homozygous samples. (a) ant.loc5– no heterozygotes; (b) *ant.loc9* – heterozygotes (n=50) cluster near 0, suggesting recessive alternate allele; (c) pale.loc4– heterozygotes show a broad intermediate distribution, consistent with incomplete dominance.; (d) serr.loc5 – no heterozygotes SNP

#### Table S1-S2

**Table S1.** Architecture of the proposed hybrid (3D–2D–1D) CNN model.

| Layer / Block | Layer Type | Kernel / Pool Size | Filters | Stride | Output Shape | Activation |
|---|---|---|---|---|---|---|
| Input | – | – | – | – | 20 × 20 × 224 | – |
| Add Channel | ExpandDims | – | – | – | 20 × 20 × 224 × 1 | – |
| 3D Block | Conv3D | 3×3×3 | 32 | 1 | 20×20×224×32 | ReLU |
| | BatchNorm | – | – | – | 20×20×224×32 | – |
| | MaxPool3D | 2×2×2 | – | 2 | 10×10×112×32 | – |
| | Dropout (0.2) | – | – | – | 10×10×112×32 | – |
| | Conv3D | 3×3×3 | 64 | 1 | 10×10×112×64 | ReLU |
| | BatchNorm | – | – | – | 10×10×112×64 | – |
| | MaxPool3D | 2×2×2 | – | 2 | 5×5×56×64 | – |
| | Dropout (0.3) | – | – | – | 5×5×56×64 | – |
| Reshape (2D) | Reshape | – | – | – | 5×5×3584 | – |
| Reshape (1D) | Reshape | – | – | – | 25×3584 | – |
| Spatial Branch | SeparableConv2D | 3×3 | 32 | 1 | 5×5×32 | ReLU |
| | BatchNorm | – | – | – | 5×5×32 | – |
| | MaxPool2D | 2×2 | – | 2 | 2×2×32 | – |
| | Dropout (0.3) | – | – | – | 2×2×32 | – |
| | SeparableConv2D | 3×3 | 64 | 1 | 2×2×64 | ReLU |
| | BatchNorm | – | – | – | 2×2×64 | – |
| | GlobalAvgPool2D | – | – | – | 64 | – |
| | Dropout (0.4) | – | – | – | 64 | – |
| Spectral Branch | Conv1D | 3 | 32 | 1 | 25×32 | ReLU |
| | BatchNorm | – | – | – | 25×32 | – |
| | MaxPool1D | 2 | – | 2 | 12×32 | – |
| | Dropout (0.3) | – | – | – | 12×32 | – |
| | Conv1D | 5 | 64 | 1 | 12×64 | ReLU |
| | GlobalAvgPool1D | – | – | – | 64 | – |
| | Dropout (0.4) | – | – | – | 64 | – |
| Fusion | Concatenate | – | – | – | 128 | – |
| | Dense | – | 128 | – | 128 | ReLU |
| | Dropout (0.5) | – | – | – | 128 | – |
| | Dense (Output) | – | 1 | – | 1 | Sigmoid |

"— indicates that the corresponding hyperparameter is not applicable to that layer type (e.g., batch normalisation has no kernel size, filters, or stride; pooling layers have no activation function)."

**Table S2.** Performance of the Randomly Selected SNPs models . Highlighted SNPs are those with high AUROC scores (>0.700)

| SNP | Accuracy | F1-Score | AUC Score |
|---|---|---|---|
| MAF_0.462_chr8_285785106 | 0.641±0.024 | 0.625±0.018 | 0.638±0.031 |
| MAF_0.411_chr2_153383894 | 0.684±0.051 | 0.662±0.025 | 0.676±0.033 |
| MAF_0.240_chr1_60823807 | 0.638±0.042 | 0.625±0.026 | 0.623±0.022 |
| MAF_0.457_chr1_157832652 | 0.649±0.033 | 0.640±0.031 | 0.644±0.019 |
| MAF_0.325_chr5_142494040 | 0.688±0.065 | 0.679±0.021 | 0.681±0.050 |
| MAF_0.323_chr1_57453753 | 0.701±0.009 | 0.684±0.065 | 0.691±0.071 |
| MAF_0.225_chr1_56267926 | 0.706±0.065 | 0.701±0.023 | 0.700±0.062 |
| MAF_0.378_chr8_67964453 | 0.709±0.044 | 0.700±0.011 | 0.701±0.033 |
| MAF_0.212_chr4_76087307 | 0.715±0.021 | 0.695±0.023 | 0.709±0.026 |
| MAF_0.223_chr2_10999703 | 0.729±0.016 | 0.701±0.020 | 0.709±0.001 |
| MAF_0.291_chr1_99864858 | 0.726±0.021 | 0.705±0.011 | 0.718±0.025 |
| MAF_0.282_chr4_273646392 | 0.659±0.032 | 0.651±0.039 | 0.657±0.010 |
| MAF_0.500_chr3_197520551 | 0.663±0.017 | 0.647±0.002 | 0.655±0.018 |
| MAF_0.263_chr8_169794280 | 0.621±0.029 | 0.608±0.026 | 0.611±0.034 |
| MAF_0.317_chr1_149106940 | 0.625±0.017 | 0.608±0.035 | 0.616±0.021 |
| MAF_0.356_chr3_194407913 | 0.642±0.009 | 0.611±0.028 | 0.618±0.040 |
| MAF_0.293_chr4_209068456 | 0.640±0.028 | 0.629±0.013 | 0.621±0.020 |
| MAF_0.452_chr2_79994966 | 0.625±0.035 | 0.625±0.019 | 0.625±0.028 |
| MAF_0.470_chr8_291297929 | 0.634±0.051 | 0.634±0.022 | 0.634±0.021 |
| MAF_0.408_chr1_69487126 | 0.667±0.039 | 0.629±0.025 | 0.637±0.011 |
| MAF_0.328_chr2_102271790 | 0.670±0.020 | 0.642±0.025 | 0.658±0.325 |
| MAF_0.305_chr7_176522372 | 0.666±0.018 | 0.658±0.007 | 0.661±0.031 |
| MAF_0.293_chr8_162260842 | 0.662±0.026 | 0.658±0.009 | 0.662±0.029 |
| MAF_0.253_chr2_25407516 | 0.669±0.015 | 0.658±0.051 | 0.662±0.017 |
| MAF_0.267_chr7_2565051 | 0.680±0.020 | 0.659±0.017 | 0.662±0.015 |
| MAF_0.202_chr1_20044862 | 0.677±0.024 | 0.661±0.008 | 0.667±0.029 |
| MAF_0.188_chr2_43528370 | 0.662±0.061 | 0.655±0.027 | 0.668±0.007 |
| MAF_0.302_chr2_39421306 | 0.683±0.018 | 0.670±0.060 | 0.677±0.021 |
| MAF_0.389_chr7_3607375 | 0.688±0.021 | 0.670±0.025 | 0.679±0.036 |
| MAF_0.487_chr8_100527853 | 0.675±0.013 | 0.659±0.018 | 0.679±0.052 |
| MAF_0.430_chr2_44898753 | 0.646±0.014 | 0.627±0.031 | 0.639±0.030 |
| MAF_0.219_chr4_161705179 | 0.651±0.053 | 0.626±0.017 | 0.639±0.044 |
| MAF_0.365_chr4_187674605 | 0.644±0.042 | 0.631±0.014 | 0.639±0.022 |
| MAF_0.322_chr3_179343248 | 0.650±0.088 | 0.641±0.022 | 0.645±0.052 |
| MAF_0.474_chr4_227639352 | 0.654±0.007 | 0.639±0.050 | 0.648±0.011 |
| MAF_0.351_chr4_130567386 | 0.655±0.016 | 0.650±0.008 | 0.653±0.003 |
| MAF_0.418_chr5_127592945 | 0.660±0.023 | 0.647±0.014 | 0.654±0.016 |
| MAF_0.186_chr2_79286655 | 0.685±0.042 | 0.670±0.044 | 0.679±0.025 |
| MAF_0.306_chr4_321098901 | 0.646±0.018 | 0.627±0.037 | 0.631±0.142 |
| MAF_0.439_chr9_52340925 | 0.590±0.053 | 0.576±0.019 | 0.587±0.013 |
| MAF_0.492_chr7_57482235 | 0.601±0.030 | 0.588±0.023 | 0.591±0.019 |
| MAF_0.419_chr4_49301152 | 0.611±0.042 | 0.585±0.091 | 0.594±0.027 |
| MAF_0.431_chr2_92080435 | 0.615±0.011 | 0.592±0.041 | 0.595±0.035 |
| MAF_0.279_chr6_15159817 | 0.593±0.029 | 0.587±0.012 | 0.596±0.016 |

| | | | |
|---|---|---|---|
| MAF_0.362_chr2_173553393 | 0.623±0.035 | 0.605±0.019 | 0.601±0.028 |
| MAF_0.386_chr4_286866607 | 0.615±0.044 | 0.609±0.016 | 0.604±0.016 |
